# Endothelial cell RpL17-dependent translational control mediates intima-media thickening in response to disturbed flow

**DOI:** 10.64898/2026.05.21.726977

**Authors:** Mary Wines-Samuelson, Sayantani Chowdhury, Sharon Senchanthisai, Michal Shaposhnikov, Mark Sowden, Bradford C. Berk

**Author notes:** These authors contributed equally to this work (Mary Wines-Samuelson, Sayantani Chowdhury). Author email addresses. Correspondence to: Bradford C. Berk, M.D., Ph.D. University of Rochester Medical Center Box URNI, 601 Elmwood Ave Rochester, NY 14642.

## Abstract

**Background:** Carotid intima-media thickening (IMT) is a major risk factor for cardiovascular disease (CVD). The large ribosomal subunit protein 17 (Rpl17) was recently reported as a CVD-associated gene; however, ribosomal mutations generally are not associated with vascular dysfunction. We have created a novel genetic model of decreased RpL17 in endothelial cells (EC) to determine how changes in endothelial ribosome expression cause IMT.

**Methods:** EC-restricted RpL17 heterozygous mice (Cdh5-Cre; *RpL17^fl/wt^,* or Rpl17-Het), were generated and subjected to sham or partial carotid ligation (PCL) surgery of the left artery to induce acute disturbed (d)-flow *in vivo*. Carotids were harvested on day 14 for quantitative tissue immunostaining. Purified mouse and human EC *in vitro* were exposed to steady (s)-flow or d-flow using cone viscometry, and collected for flow cytometry, protein expression, electron microscopy, or purification of ribosomes. Human carotid samples from healthy and endarterectomy patients were used for tissue analysis.

**Results:** Carotids from RpL17-Het mice with PCL-induced d-flow showed increased IMT relative to RpL17-WT controls. In addition, RpL17 protein levels were decreased in regions of d-flow compared to s-flow. Increased levels of ER stress markers were observed by carotid immunostaining, as well as activation of the integrated stress response (ISR) in RpL17-Het EC. Analysis of mRNAs bound to polysomes vs. monosomes in EC-RpL17-Het revealed increased translational efficiency of key regulators of glycolysis, redox, inflammation, matrix, and endothelial-to-mesenchymal transition (EndMT). Metabolic profiling by Seahorse assay showed enhanced anaerobic glycolysis and decreased oxidative respiration in RpL17-Het EC, consistent with the translational efficiency data. Immunostaining of carotids identified upregulated EC inflammation and EndMT.

**Conclusions:** Our data support RpL17 as a key mediator of EC phenotypic modulation that causes IMT in response to d-flow. We show a novel pathway for d-flow-mediated IMT: endoplasmic reticulum stress and activation of the ISR. These changes alter translational efficiency and reprogram EC cell cycle, metabolism, and redox state in the presence of d-flow to cause IMT, a precursor to cardiovascular pathology.

## Introduction

Intima-media thickening (IMT) of the vascular wall is widely accepted as a susceptibility factor for the development of atherosclerosis. In fact, carotid IMT is used as a diagnostic measurement to evaluate atherosclerosis in the carotid. Carotid atherosclerosis is a potentially fatal disease, in that plaque rupture can cause a stroke. Although much is known regarding the contribution of high-fat diet and lipids to vascular dysfunction, less is understood regarding genetic risk factors which underlie vulnerability to development of carotid IMT and atherosclerosis. Endothelial cells (EC), which form the innermost cellular layer within blood vessels, contain mechanosensors to detect changes in blood flow. Specifically, in response to slow, turbulent and disturbed (d-flow), EC express receptors for inflammatory cell binding and activation, as well as changes in their own function. Their normal quiescent state is disrupted by d-flow as occurs in vessels narrowed by atherosclerotic plaque. Under chronic conditions of exposure to high glucose or cholesterol (e.g., diabetes or hyperlipidemia), EC are “activated” to pro-inflammatory, pro-migratory programs and decreased vasodilator function (Fig. 1). With EC activation, the cells can enter the cell cycle or undergo endothelial to mesenchymal transition (EndMT) to contribute to IMT and vascular remodeling.

**Figure 1:**
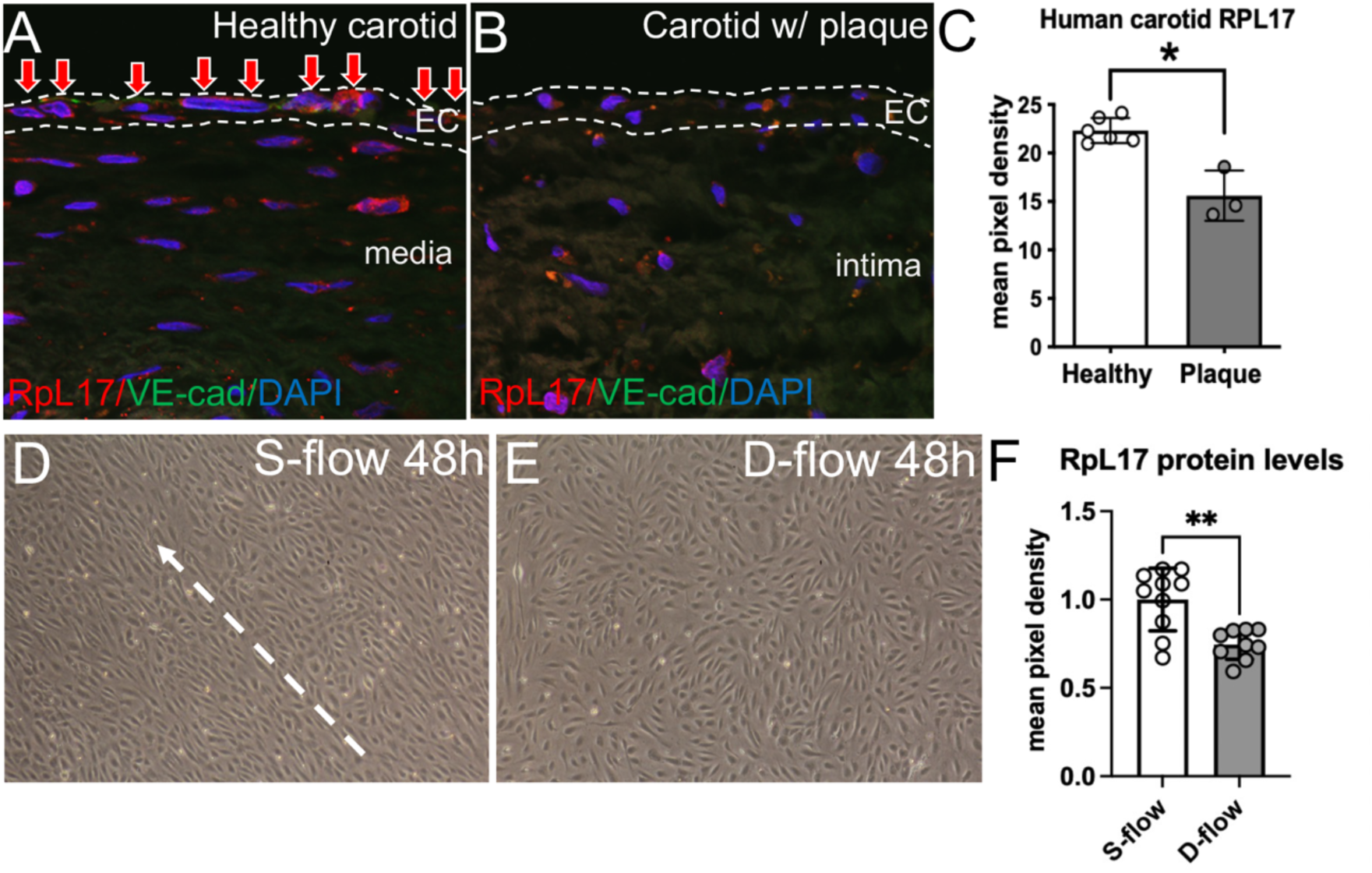
Human EC in carotid arteries with plaque or cultured with disturbed (D) flow have decreased RPL17 levels. (A-C) Human carotid sections immunostained for RpL17 in plaque-bearing vessel (B) & healthy vessel EC (A, red arrows), quantified in (C). (D-F) Human umbilical vein EC (or HUVECs) cultured in steady (S) flow (panel D) or disturbed (D) flow (panel E); protein lysates from HUVECs assayed for RPL17 levels quantified in (F). *EC, endothelial cell; *, p < 0.05; **, p < 0.01*.

Previous genetic studies from our laboratory identified three key chromosomal loci, *Intima modifier-1,2,* and *3,* which conferred IMT susceptibility following partial carotid ligation (PCL) of inbred strains of mice in the absence of other atherogenic factors2. Within the *Im-3* locus, one gene encoding the large ribosomal subunit protein 17 (RpL17), exhibited very low expression in strains with the highest levels of IMT. A recent coronary artery disease (CAD) GWAS study identified 64 novel genetic loci as putative candidates for CAD; SNPs in RPL17 were found to be significantly associated with vascular disease in humans^1^. Only one other ribosomal protein, a mitochondrial protein MRPL14, was similarly identified in the study. Decreased expression of a ribosome protein would be expected to inhibit protein translation and cell growth. Surprisingly, while there was a 50% decrease in global protein translation, there was a very significant increase in cell growth. We discovered that activation of the integrated stress response (ISR) and Atf4, led to significant increases in translational efficiency of a special group of mRNAs. These included mRNAs with putative cis-acting regulatory elements such as 5’ upstream open reading frames (uORFs), internal ribosome entry sites (IRES), and binding sites for several microRNAs (miRNAs) and RNA binding proteins. Here we show the mechanisms by which these changes in translational efficiency and endothelial phenotype promote cell growth, EndMT and IMT. Our data support a novel pathway for d-flow-mediated IMT via ISR activation that reprograms the translational state of EC. Flow pattern plays a major role in regulating the progression to IMT, with d-flow stimulating EC inflammation, proliferation, and EndMT.

## Materials and Methods

### Data availability

Detailed information regarding the data, methods, and materials used for this study are provided in the online Supplemental Material.

### Mice

Mice were maintained under a standard 12-hour light-dark cycle with regular chow and water *ad libitum* in accordance with institutional UCAR standards. The floxed RpL17 mice were generated by creating founder chimeras with targeted heterozygous p5 male JM8 C57BL/6N embryonic stem cells (ESCs) purchased from the European Mouse Mutant Cell Repository (EuMMCR) as part of EUCOMM (MGI Allele ID# 2448270, Clone IDs HEPD0801_5_E04 and HEPD0801_5_H03, allele name *Rpl17^tm3a(EUCOMM)Hmgu^*; ^1^). ESCs were expanded and injected into blastocysts by the University of Rochester Medical Center Transgenic Facility to generate founder mice. Floxed RpL17 heterozygotes were maintained on a C57 BL/6 background, and bred to Cdh5-Cre (stock #017968, strain B6;129-Tg(Cdh5-cre)1Spe/J from Jackson Labs) transgenic mice^2^.

### Translational efficiency analysis

Luciferase mRNA was spiked into each fraction as a normalization control for RNA content. An equal volume (7.5 µL) of purified total RNA (range: 90-430 ng/µL) was reverse transcribed using iScript cDNA synthesis (Bio-Rad). Quantitative real-time PCR amplification of transcripts was performed using 4 µL diluted cDNA (range: 13-64 ng RNA equivalents of cDNA), 1 µM specific primers (see supplemental methods), and SybrGreen PCR reagent (Bio-Rad). Cycle parameters were a 2-step protocol of 95°C denaturation and 60°C annealing/extension for 40 cycles. Luciferase amplification served as an internal control.

### Statistical analysis

Statistical analysis was performed using GraphPad Prism 9. Quantitative data was collected by an experimenter blinded to the genotype or treatment of sample. For fluorescent image analysis, Fiji/ImageJ2 (v.2.3.0) image processing software (NIH) was used on single-channel images with the same threshold applied per experiment, and mean pixel density was recorded per image. Unpaired 2-tailed Student *t*-test with Welch’s correction was used when comparing two groups. A value of *p* < 0.05 was considered statistically significant. the number of * indicates the following statistical significance *p<0.05, **p<0.01, ***p<0.005, and ****p<0.001. Replicates are biological and indicate number of mice used, except for subcellular structures (e.g., ribosome particle density), in which case the number of fields examined is equal to *n.* Error bars on graphs indicate the standard deviation of the sample group.

## Results

### RPL17 is decreased in EC in human carotids exposed to d-flow and atherosclerosis

RpL17 was initially discovered as a candidate gene for the *Intima modifier-3 (Im3)* locus on mouse chromosome 18 ^3^. A subsequent study by the Berk laboratory in 2012 performed RpL17 siRNA knockdown of all cells in the mouse carotid artery by pluronic gel implantation, and observed increased intima thickness with knockdown ^4^. In the same study, cultured rat aortic smooth muscle cells showed increased proliferation after siRpL17 knockdown ^4^. However, these data did not address the role of RpL17 in endothelial cells, an essential immune and flow signal modulator for vascular growth responses. In addition, it was not known whether human RPL17 vascular function would be conserved from mice to humans. To determine whether RpL17 levels could be altered in endothelial cells exposed to disturbed flow in diseased human carotid tissue plaques, we examined endarterectomy samples by immunostaining for RpL17 (Fig. 1). Carotids were prepared for analysis by orienting with respect to the bifurcation at defined distances along the rostral-caudal axis (see Methods). RPL17 was expressed at moderate to high levels in EC in the healthy carotid (Fig. 1A, red arrows), and in medial cells. However, in plaque-bearing carotids, EC and medial cells contained significantly reduced levels of RpL17 (Fig. 1B vs. 1A, quantified in Fig. 1C).

To confirm our results and evaluate whether the effects on RpL17 expression were due primarily to disturbed flow and not other pathologies associated with vascular plaque, we cultured human umbilical vein EC (HUVEC) cells in the presence of steady (s) flow or disturbed (d) flow and quantified levels of RPL17 protein from the cells by western blotting (Fig. 1D-F). In the presence of d-flow, RPL17 levels decreased by 25% relative to HUVEC exposed to s-flow (Fig. 1F; p <0.01), thus supporting our immunostaining data from human endarterectomy EC that revealed ribosomal RPL17 is a flow-responsive protein. Moreover, although RpL17 was a candidate gene that mapped within the *Intima modifier-3* genomic region of mouse chromosome 18, these results were the first demonstration that endothelial RpL17 itself was a flow-responsive protein in humans.

### RpL17 mRNA and protein are reduced in Cdh5-Cre; RpL17^fl/wt^ EC

To functionally assess the role of RpL17 in vascular EC *in vivo*, we generated an EC-specific, Cdh5-Cre; RpL17 heterozygous mouse (hereafter referred to as EC-RpL17 het). Specifically, we bred RpL17^fl/fl^ mice containing LoxP sites flanking exons 6 and 7 of RpL17 with mice carrying the vascular endothelial specific cadherin-5-Cre (VE-cadherin, or Cdh5-Cre) transgene, whose expression is limited to EC ^5^. We found that complete loss of Rpl17 in EC was embryonic lethal (Berk lab, unpublished data), which is typical of many ribosomal protein homozygous knockouts ^6^. In contrast, EC-RpL17-Het mice live to adulthood with no apparent phenotype (Berk lab, unpublished data). Both males and females have normal body weights, fertility, and survival after PCL (data not shown).

In order to determine whether the RpL17-Het mice had reduced levels of RpL17, the thoracic aorta was harvested from 6-week-old WT and RpL17-Het mice and subjected to *en face* staining (Fig. S1A-C). RpL17-Het aortas showed significantly lower immunostaining levels of RpL17 compared to their WT counterparts (Fig. S1B vs. S1A, S1C), confirming the RpL17 genetic knockdown in mice. Further, western analysis of RpL17 knockdown in ribosome-enriched lysates from mouse lung microvascular ECs (or MLMEC) from RpL17-Het vs. controls revealed that RpL17 protein was present but reduced 51% in RpL17-Het EC (Fig. S1D-F, S1H). In contrast, levels of another 60S ribosomal exit tunnel protein, RpL35, showed no significant change in RpL17-Het EC ribosomes (86% of RpL17-WT, p> 0.05; Fig.S1H), as well as a 40S ribosomal protein, RpS19 (114%, p> 0.05). These data demonstrated that we established a suitable RpL17 knockdown EC model system *in vitro* to study the mechanisms and functional consequences of decreased RpL17 in EC.

### Effect of flow pattern on RpL17 mRNA and protein levels in EC

To establish whether RpL17 would respond to differential flow status in mouse EC, we performed RpL17 immunostaining on mouse carotids with or without PCL (Fig. S1I-L). In RpL17-HET versus RpL17-WT sham-operated carotids (Fig. S1J vs S1I; quantitated in Fig. S1M), RpL17 immunostaining was significantly reduced by a small amount in the EC layer (12±2.2%). Rpl17 immunostaining in carotid EC exposed to d-flow after PCL showed a very significant reduction in RpL17 in the EC layer (23±3.5%; Fig. S1L vs. S1K; quantitated in Fig. S1M). Taken together, these results identified RpL17 as flow-sensitive in mouse EC *in vivo*.

### RpL17-Het carotids have increased intima-media thickening (IMT) after PCL

Quantitative analysis of carotid sections collected 14 days after PCL revealed that EC-RpL17-Het male carotids had increased intimal thickening of the ligated (left) carotid compared to the ligated RpL17-WT carotid, as measured by compartment volume (Fig. 2A-C, Fig. 2E). Note that these experiments were performed with C57/BL6 mice that have very little IMT in the ligated carotid ^7^. To further examine changes in vascular response to d-flow, we measured the volume of the external elastic lamina (EEL), defined as the volume between the media and adventitia. Consistent with the increase in intimal thickness, we observed increased EEL volume in EC-RpL17-Het carotids (p<0.01; Fig.2G). Notably, the lumen of EC-RpL17-Het male carotids was significantly dilated with respect to RpL17-WT carotids (Fig. 2D). Thus, decreased levels of RpL17 in EC uniquely alter male vascular remodeling via growth of the intima and EEL compartments. Since IMT was only present in male EC-RpL17-Het mice, males were used for the majority of experiments.

**Figure 2:**
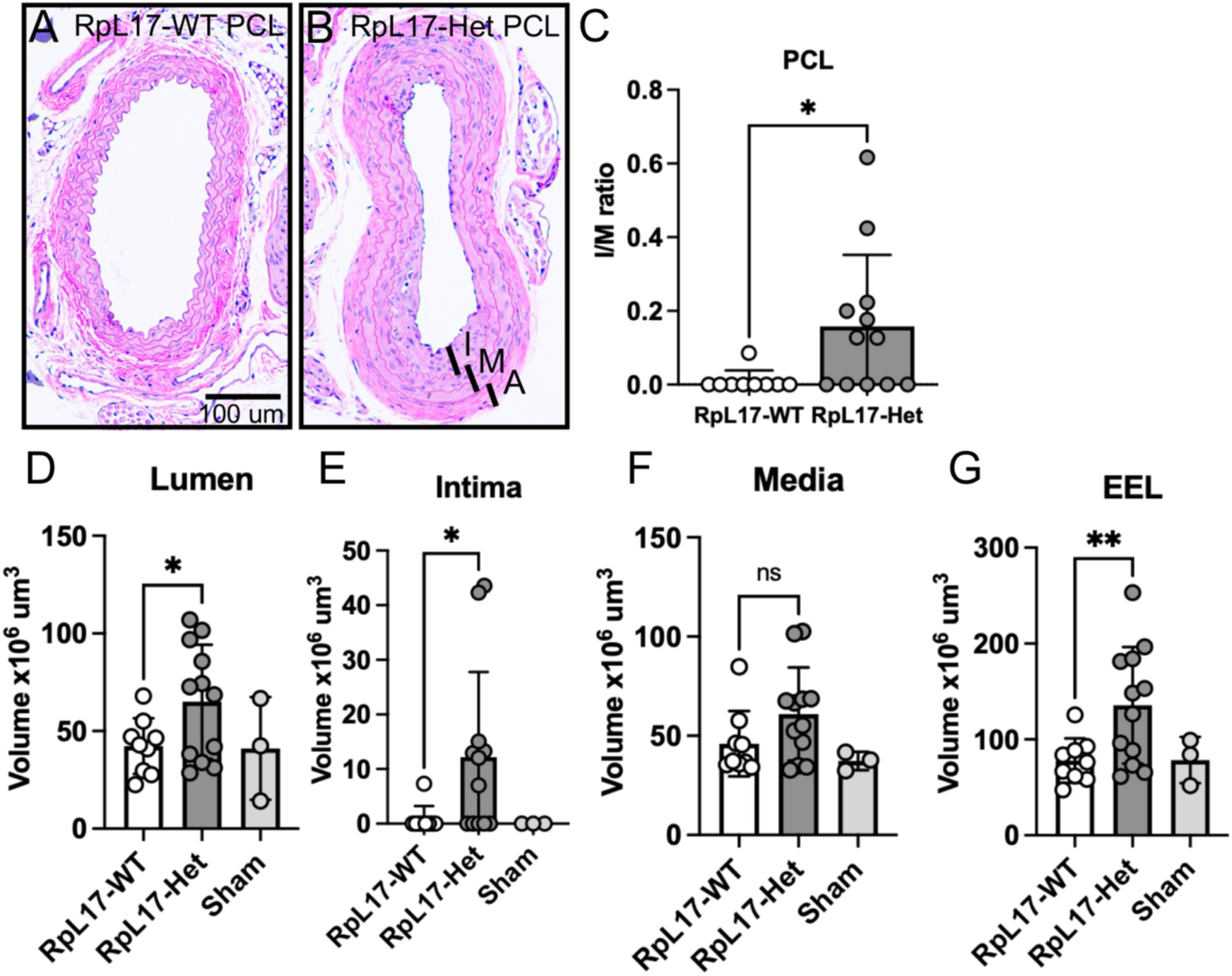
Reduction of RpL17 in carotid EC promotes I-M thickening (IMT) after partial ligation. (A,B) Images of H&E-stained carotid sections taken from male mice at 14 days post-partial ligation. (C-G) Morphometric analysis of ligated (PCL) carotids from males. Increased levels of remodeling were measured in EC-RpL17 vs. Control by measuring the I/M ratio (C), the lumen (D), the intima (E), and the external elastic lamina (EEL, G). *NS, not significant (p> 0.05); PCL, partial carotid ligation; I, intima; M, media. * indicates p<0.05; ** p<0.01; and *** p<0.001*.

### RpL17-Het EC have increased proliferation, similar to RpL17-Het EC exposed to d-flow

It has been verified that d-flow is a major stimulus for growth in the vessel wall as shown by IMT and atherosclerosis ^8–10^. Previously, RpL17 was identified as an intima modifier candidate gene and subsequently shown to promote proliferation of vascular smooth muscle cells when reduced by RpL17 siRNA treatment ^4^. Therefore, we hypothesized that decreased RpL17 in RpL17+/- EC might stimulate re-entry of normally quiescent EC into the cell cycle and sustained cell proliferation. To test our hypothesis, we used cultured MLMEC from RpL17-WT and RpL17-Het lungs, and then measured expression of cyclin D1 (Fig. 3A-B). Strikingly, RpL17-Het MLMEC revealed a 3.8-fold increase in the level of cyclin D1 relative to RpL17-WT MLMEC (Fig. 3B), suggesting dysregulation of the G1-to-S transition with reduced levels of RpL17. To quantitively assess the EC cell cycle, we analyzed cultured MLMEC by cell sorting on the basis of nuclear DNA content. Flow cytometry (FACS) analysis of RpL17+/- EC vs. RpL17+/+ EC showed a 2-fold increase in the number of RpL17+/- EC in S-phase and M-phase of the cell cycle (Fig. 3C). We next studied the effects of exposure to d-flow on EC proliferation using the PCL model. We immunostained RpL17-WT and RpL17-HET carotids 14 d after PCL. There was a 150% increase in proliferating cell nuclear antigen (PCNA)-positive cells in RpL17HET vs. RpL17-WT carotids (Fig. 3E/E’ vs. 3D/D’; quantitated in Fig. 3F). Importantly, PCNA levels were normal in RpL17-HET sham carotids (data not shown). Together, these results demonstrate that reduced RpL17 in EC stimulates proliferation.

**Figure 3:**
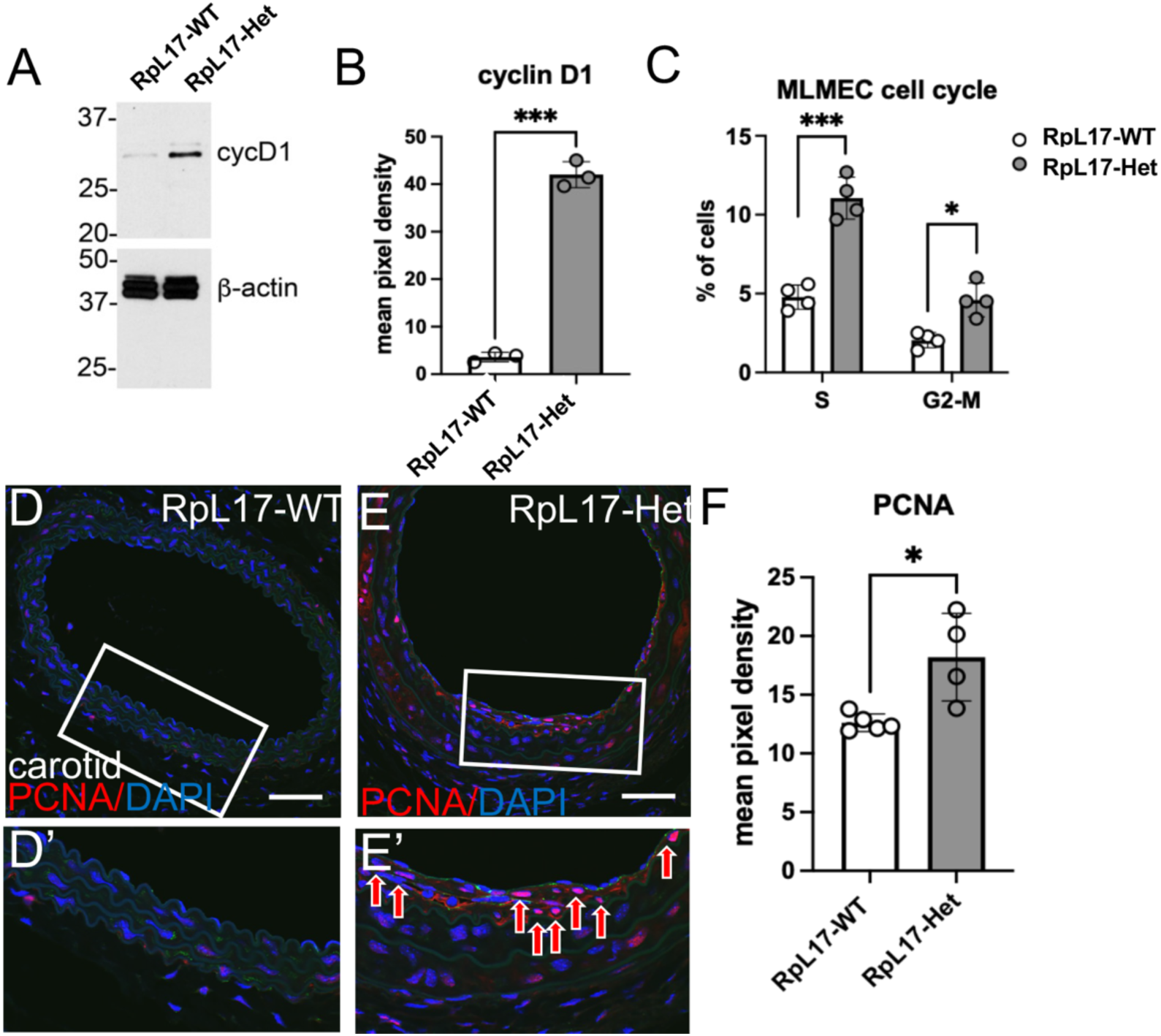
RpL17+/- EC are proliferative. (A-C) Cultured MLMEC from RpL17-WT and RpL17-Het lungs. (A), Cyclin D1 (cycD1) western blots (quantified in (B)). (C), cell sorting analysis on nuclear DNA content to determine % of cells in S- and M-phase of the cell cycle. (D,E) Immunostaining of mouse carotids with antibody against proliferative marker PCNA (quantified in F). *\*, p< 0.05; ***, p< 0.001*.

### EndMT is increased in EC-RpL17-Het carotids after PCL

In addition to increased proliferation of ECs in RpL17-Het cultured MLMEC and in EC-RpL17-Het carotids after PCL, we reasoned that the increased IMT phenotype of EC-RpL17-Het mice may also contain cells undergoing EndMT, since previous scRNA-seq data from the Jo lab using the PCL model showed a tight correlation between d-flow (EC stress exposure) and increases in EC fate switching to EndMT ^11^. To test the hypothesis, we immunostained PCL carotids for the EndMT transcription factor Snail (Fig. 4A-B; quantified in Fig. 5C), and the TGF-b signal transduction factor phospho-Smad2 (Fig. 4D-E; quantitated in Fig. 4F), which are expressed by EC as they undergo EndMT. RpL17-WT carotids after PCL showed little intima, and markedly reduced Snail, especially in the media (Fig. 4A/A’). However, RpL17-Het carotids after PCL showed significant increases in the intima and media of Snail (155%, p< 0.05, Fig. 4B/B’ vs. 4A/A’, 4I) and phospho-Smad2 (360%, p<0.01; Fig. 4E/E’ vs. 4D/D’, 4J), as well as increased smooth muscle alpha actin in the intima (data not shown). In summary, the finding of phospho-Smad and Snail in the media strongly support the concept that a significant number of cells have undergone EndMT.

**Figure 4:**
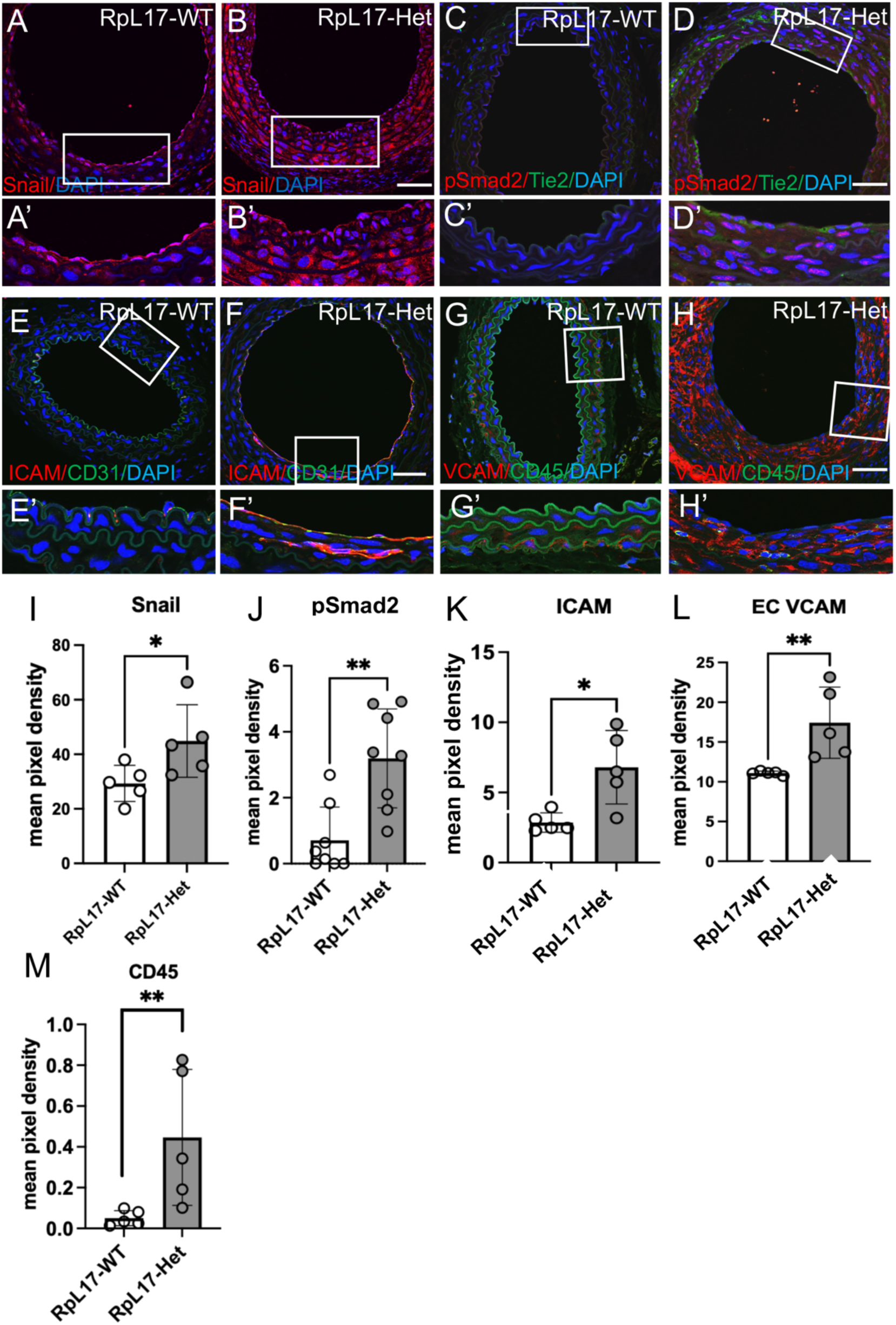
Increased EndMT and EC inflammation in RpL17-Het ligated carotids. (A,B) Immunostaining of ligated carotids with antibody against EndMT transcription factor Snail (quantified in E; p< 0.05). (C,D) Activation of the EndMT-associated TGF-beta transcription factor phospho-Smad2 in carotids from RpL17-WT (C,C’) vs. RpL17-Het (D,D’) (quantified in I,J) revealed that both markers Snail and pSmad2 are increased in RpL17-Het carotids after PCL. Scalebar, 50 microns. (E-H) Activation of adhesion molecules ICAM-1 and VCAM-1 in carotids from RpL17-WT (E,G) and E’,G’) vs. RpL17-Het (F,H) and (F’,H’) respectively; quantified in (K.L). (M) Quantification of CD45-positive monocyte-derived inflammatory cells detected in G and H revealed increased CD45+ cells in Rpl17-Het carotids (green cells in H’ vs. G’). Scalebar, 50 microns.

**Figure 5:**
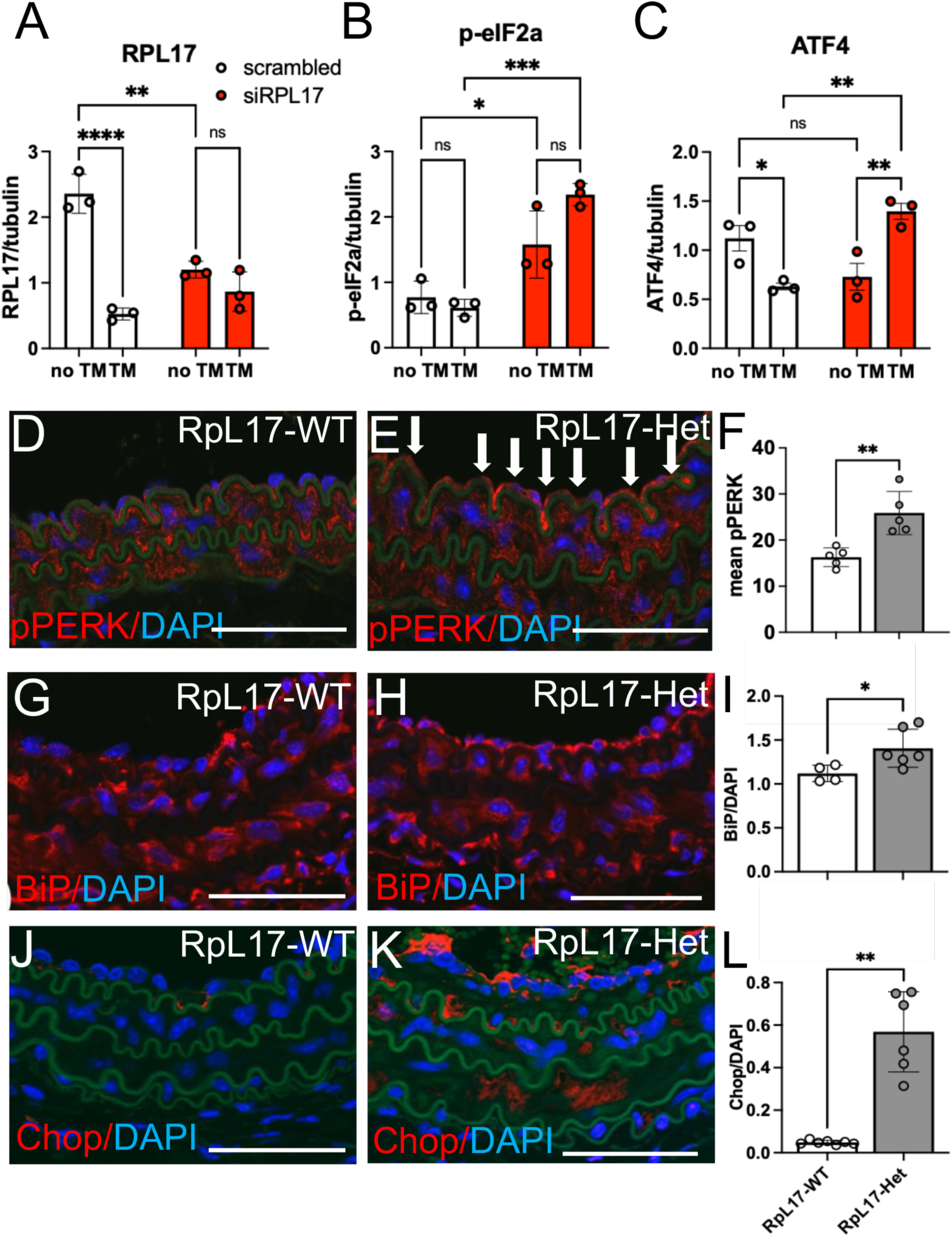
Activation of ISR and ER stress in EC-RpL17. Quantification of immunoblots from untreated (“no TM”) or tunicamycin-treated (TM) HUVECs transfected with either RPL17+/+ (scrambled) or siRNA against RPL17 (siRPL17) probed for RPL17 (A), p-eIF2α (B), and ATF4 (C). siRPL17 EC showed increased expression of p-eIF2alpha and Atf4, indicating activation of the ISR, which attenuates global translation. (D-L) Immunostaining of PCL carotid sections from RpL17-WT and RpL17-Het mice with antibodies specific for stress kinase p-PERK (D-E, quantified in F), and markers of ER stress BiP (G-H, quantified in I) and Chop (J-K, quantified in L). Scalebars, 50 microns.

### Inflammation is upregulated in EC-RpL17 carotids after PCL

Expression of pro-inflammatory cell surface proteins on EC and recruitment of immune cells are early steps in vascular remodeling ^12–14^. Therefore, we immunostained PCL vessels for ICAM1, VCAM1, and CD45 (Fig.4). In RpL17-Het vs. RpL17-WT PCL carotids, we found increases of 3-fold in ICAM1 (Fig.4F/F’ vs. 4E/E’, quantified in Fig. 4J); 1.6-fold in VCAM1 (Fig. 4H/H’ vs. 4G/G’, quantified in Fig. 4L); and 9-fold in CD45+ leukocytes (Fig. 4H/H’ vs. 4G/G’, quantified in Fig. 4M). There were minimal amounts of all three inflammation-associated markers staining in sham vessels (data not shown). Thus, decreased RpL17 in EC increases vascular inflammation after PCL.

### RpL17-Het carotid EC have increased oxidative stress after PCL

Since pro-inflammatory immune cells are a major source of ROS in inflamed tissues, and RpL17-Het carotids show increased EC inflammation and CD45+ leukocytes, we investigated whether the RpL17-Het carotids would have increased oxidative damage after PCL. Immunostaining of PCL carotid sections from RpL17-Het vs. RpL17-WT mice with anti-4-hydroxynonenal (4HNE) revealed increased lipid peroxidation (Fig. S2B vs. S2A, quantitated in Fig. S2E), and immunostaining with anti-8-oxo-2’-deoxyguanosine (8-oxo-dG) revealed increased DNA oxidation levels (Fig. S2D vs. S2C, quantitated in Fig. S2F). Together, these results identify a decreased ability of RpL17-Het EC to maintain redox homeostasis in inflamed vessels exposed to d-flow.

### Effect of RpL17 haploinsufficiency on protein expression and the ISR

In light of the broad range of effects we observed in EC-RpL17-Het carotids exposed to d-flow (matrix remodeling, EC inflammation and proliferation, and EndMT), we reasoned that there might be a global change in the status of translation capacity of EC ribosomes with a 50% reduction in RpL17. To address this possibility, we treated RpL17 HUVEC with two RpL17 siRNAs (RpL17-siB and -siC), which reduced RPL17 protein by ∼60% after 48 hours of treatment (Fig. S3B-C). siRPL17-HUVEC were exposed to puromycin, a charged tRNA analog, to label all actively translating proteins, and their lysates analyzed by western blotting with anti-puromycin antibody (Fig. S3D-E). We found a ∼35% reduction in puromycylation, and a 35% reduction by immunostaining (data not shown) in siRpL17-treated HUVEC relative to control (scrambled siRNA)-treated HUVEC (Fig. S3D, quantitated in Fig. S3E). Thus, overall translation was significantly inhibited by decreased levels of RpL17, perhaps due to activation of the integrated stress response, or ISR.

### Decreased translation, ISR and ER stress are present in RpL17-Het EC

After confirming that EC with decreased RpL17 show attenuated global translation by puromycylation, we hypothesized that the overall decrease in translation in RpL17-Het EC could be associated with activation of the ISR pathway. The ISR conserves cellular energy when a severe stress occurs by preventing cap-dependent translation via eIF2 alpha. Phosphorylation of eIF2 alpha by stress-activated kinases (e.g. ER stress-dependent PKR-like ER kinase (PERK) impairs its ability to initiate translation within the ternary complex ^15^. We observed increased phosphorylation levels of both eIF2 alpha (Ser51; 1.8-fold) and its kinase PERK (1.7-fold) by western blot (Fig. 5A-B) and by immunohistochemistry (IHC) of PCL carotids (Fig. 5D-E). To confirm activation of the ISR, we measured levels of the downstream ISR target, Atf4. There was a 1.7±0.3-fold increase in Atf4 protein levels in RpL17+/- EC vs. RpL17+/- EC (Fig. 5A). Taken together, these data show that decreased RpL17 stimulates ISR activation in EC under static or d-flow.

With activation of the ER-resident stress kinase PERK, we investigated whether its inducer, ER stress, was activated in RpL17-Het carotid EC after PCL. Immunostaining of carotid sections for ER stress chaperone Binding immunoglobulin Protein (BiP) (Fig. 5G-I) and the chronic ER stress/unfolded protein response (UPR) transcription factor Chop (CCAAT/-enhancer-binding protein homologous protein; Fig. 5J-L) revealed significantly increased BiP-positive (1.3±0.5-fold increase; Fig. 5I) and Chop-positive (∼10-fold increase; Fig. 5L) immunoreactive cells in the RpL17-Het vs. RpL17-WT EC layer. Together, RpL17-Het versus RpL17-WT carotids had dramatically increased BiP- and Chop-immunoreactivity, clearly indicating the presence of ER stress with low RpL17 in vessels under d-flow.

To confirm ER stress, we investigated the rough ER (RER) ultrastructure of EC, since the RER contains bound ribosomes, and its area has been shown by others to enlarge in the presence of chronic ER stress and activation of the UPR^16^. We measured the dimensions of individual RER sacs by electron microscopy (EM) in RpL17-WT and RpL17-Het EC (Fig. S4) ^17^. There was increased area and width of the RER in RpL17+/-EC (red arrowheads, Fig. S4B vs. S4A). Quantification of both maximal width of the RER sacs (Fig. S4C) and area of the sacs (Fig. S4D) confirmed increases in both measures (max width: 0.062 vs. 0.045 um; p< 0.0001; area: 0.06 vs. 0.02 um^2^; p< 0.0001). In contrast, another plasma membrane structure, caveolae, were not altered in RpL17+/- EC (1.7 vs. 1.5 um^2^, p> 0.05; data not shown). These results support our hypothesis that reduced RpL17 promotes protein misfolding, ER stress, and the ISR.

### RpL17-Het EC show large changes in translational efficiency

Initially, we performed RNA-Seq with RpL17-Het and RpL17-WT EC to determine downstream effects of RpL17 reduction (data not shown); however, we observed that the majority of differentially-expressed genes (DEGs) were downregulated and had changes less than 2-fold. This suggested to us that the dominant changes in RpL17-Het EC would be at the level of translation, not transcription. Therefore, we took a targeted approach to identify pathways that would be altered post-transcriptionally to promote EndMT and IMT as we had observed in RpL17-Het carotids under d-flow. To address this, we measured translational efficiency, or the proportion of a specific mRNA bound to multiple ribosomes (in polysome fractions) vs. a single ribosome (in monosome fractions), of candidate genes known to be highly dysregulated in EC undergoing EndMT ^18,19^ (Fig. 6).

**Figure 6:**
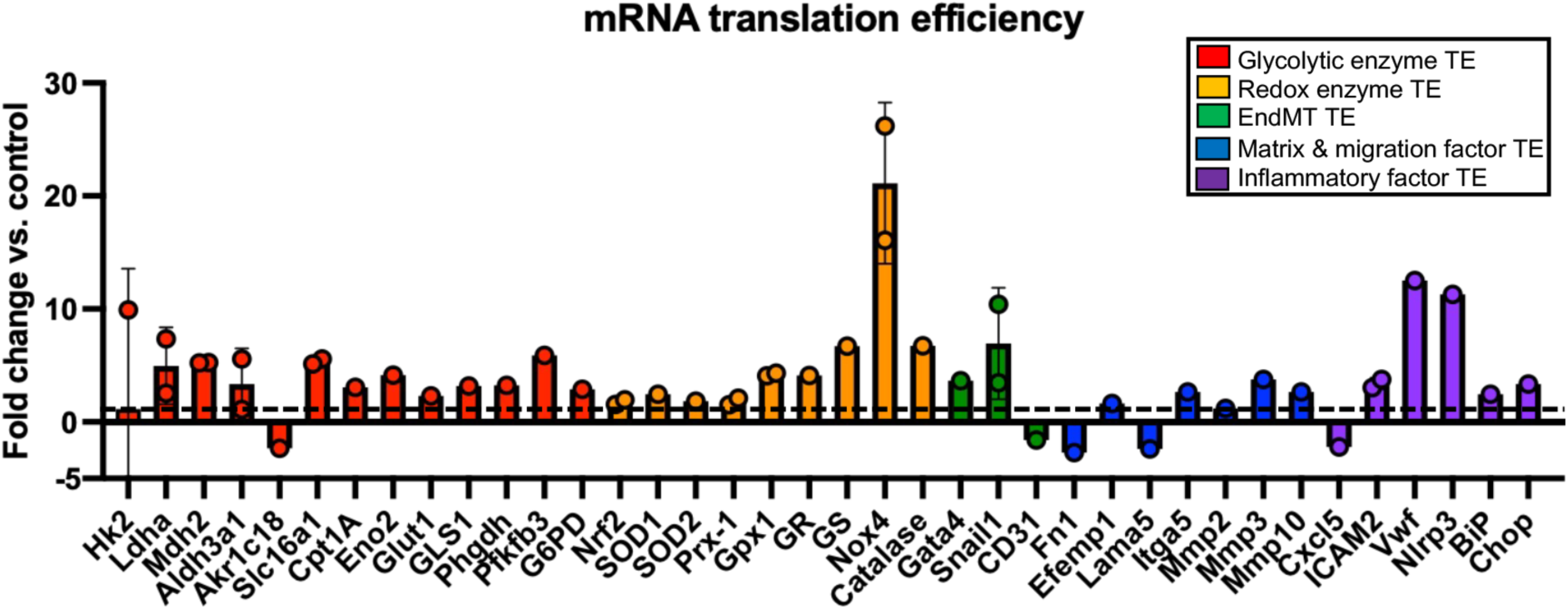
Translation efficiency (TE) measurement by qPCR of purified ribosomal fractions from MLMEC mRNA. Increased TE in EC-RpL17 vs. Control polysomes for antioxidant enzymes (Nrf2, nuclear factor erythroid 2-related factor 2; Gpx1, glutathione peroxidase 1; Prx1, peroxiredoxin 1; catalase; SOD1 and 2, superoxide dismutases 1 & 2) including a pro-oxidative, ROS-generating enzyme NADPH oxidase 4 (Nox4) and glycolysis regulatory enzymes (Ldha, lactate dehydrogenase-a; Pfkfb3, 6-phosphofructo-2-kinase/-fructose-2,6-biphosphatase 3; Slc16a1, solute carrier 16a1 (also monocarboxylate transporter-1 (MCT1), or lactate transporter); Ldha, lactate dehydrogenase-a; Glut1, glucose transporter 1; Eno2, enolase 2; Phgdh, phospho-glycerate dehydrogenase), various metabolic regulators (Mdh2, malate dehydrogenase 2; Cpt1A, carnitine palmitoyltransferase 1A; GR, glutathione reductase; and GLS1, glutaminase 1), and ER stress regulators BiP (Grp78) and Chop (CCAAT-enhancer protein binding matrix metallo-proteases Mmp3 and Mmp10, pro-inflammatory ICAM, Nlrp3, and von Willebrand factor (Vwf); in endothelial-to-mesenchymal transition (EndoMT) transcription factors Snail1 and Gata4; and in ER stress factors Chop and BiP. Decreased polysome mRNA levels associated with matrix components (fibronectin, Fn1; laminin Lama5), and cytokine receptor Cxcl5. TE represents polysome vs. monosome mRNA fold enrichment, normalized to an internal spiked-in control luciferase mRNA; polysome & monosome fractions were identified and pooled based on ribosome profiling data using equal volumes of mRNA from individual fractions. Enrichment was calculated as the fraction of the total for each mRNA across all ribosomal fractions normalized to a ribosome-free fraction, then EC-RpL17 fold change compared to Control.

We performed translational profiling of the EC by comparing amplification levels of highly-translating polysome-associated mRNAs vs. low-translating monosome-associated mRNAs extracted from purified ribosomal fractions ^20,21^. Interestingly, there was a wide range of translational efficiency of the mRNAs represented by the six functional categories found in the purified ribosomal fractions, suggesting dynamic regulation of these mRNAs (Fig. 6). A noticeable difference was observed between translational efficiency and total mRNA levels (Fig. 6 and data not shown). We also observed changes in EndMT: Gata4 and Snail1, EndMT transcription factors, showed 3-and 7-fold increases, respectively; while platelet endothelial cell adhesion molecule (PECAM-1), also known as cluster of differentiation 31 (CD31), a marker of differentiated EC, was significantly decreased (Fig. 6). Consistent with increased levels of migration and loss of EC adhesion in RpL17-Het EC, we identified reduced translational efficiency of extracellular matrix factors fibronectin and laminin, while translational efficiency of pro-migratory Mmp3 and Mmp10 were increased significantly (Fig. 6). The results suggest that RpL17-Het EC will degrade matrix and display enhanced migration, consistent with phenotypic modulation (EndMT) and vascular remodeling. Altogether, these changes likely reflect a coordinated metabolic response to promote survival that results in EC phenotypic modulation toward cells with EndMT- and inflammatory-like properties ^22,23^.

### Glycolytic metabolism is upregulated in RpL17+/- EC in vitro and in vivo

Due to the increased translational efficiency of a number of glycolytic enzymes in RpL17-Het EC vs. WT control EC (Fig. 6), we assayed the EC for glycolytic flux by Seahorse assay (Fig. S5). Extracellular acidification rate (ECAR) essentially measures the efflux of lactate, a metabolite produced and secreted during glycolysis ^24–27^. Increased basal and maximal ECAR in RpL17-Het vs. WT EC (Fig. S5A-B) was consistent with increased lactate production (Fig.S5C), and glycolysis ^24–27^. Secretion of lactate is consistent with a pro-angiogenic phenotype (proliferative and migratory), and with EndMT (Fig.3, Fig. 4). Immunofluorescence staining showed increased expression of Pfkfb3 in RpL17-Het carotids compared to RpL17-WT (Fig. S5E vs. S5D, quantified in Fig. S5F). Pfkfb3 catalyzes synthesis and degradation of fructose 2,6-bisphosphate, a rate limiting step in glycolysis. It also regulates cyclin-dependent kinase 1, thereby linking glycolysis to cell proliferation ^27–29^.

### Mitochondrial respiration and OXPHOS are impaired in RpL17-Het EC

We hypothesized that RpL17-Het EC would display functional deficits in mitochondrial respiration due to the increased glycolysis levels we had observed. Mitochondrial respiration of MLMEC was determined by Seahorse extracellular flux analyzer to assess oxygen consumption rate (OCR) and the respiratory reserve capacity, which measures the potential to increase respiration after uncoupling electron transport ^30,31^. RpL17-Het MLMEC showed 16% (p<0.05) reduced maximal levels of respiration compared to RpL17-WT EC (Fig. S6A-B). Western blot analysis of OXPHOS complex proteins showed significantly decreased levels of proteins associated with Complexes I, III, and IV, consistent with our RNA-seq analysis of downregulated genes in RpL17-Het EC (data not shown), but not Complex II or V (Fig. S6C-D). Thus, the decrease in OXPHOS-dependent respiration is likely due to decreased expression of Complex I, III, and IV subunit proteins.

### Human aortic EC require RPL17 to suppress ER stress, inflammation, and proliferation

To test whether RpL17 function is conserved between mice and humans, we examined levels of ER stress, ISR, and proliferation markers in human primary aortic endothelial cells (HAEC) treated with siRNA against RPL17 (Fig. 7). Two siRNAs directed against RPL17 (RPL17-siB and RPL17-siC) were found to reduce levels of RpL17 protein in HAEC to ∼30% of controls (scrambled siRNA-treated) after 48 hr (Fig. 7). Treatment of HAEC with RpL17-siB compared to RpL17-scrambled for 24 hr increased key markers of stress, such as BiP/Grp78 (3-fold); ICAM1 (2.5-fold); and cyclin D1 (2.5-fold) (p< 0.05; Fig. 7). These results were consistent with data from RpL17-Het MLMEC, demonstrating highly conserved function of RpL17 in EC.

**Figure 7:**
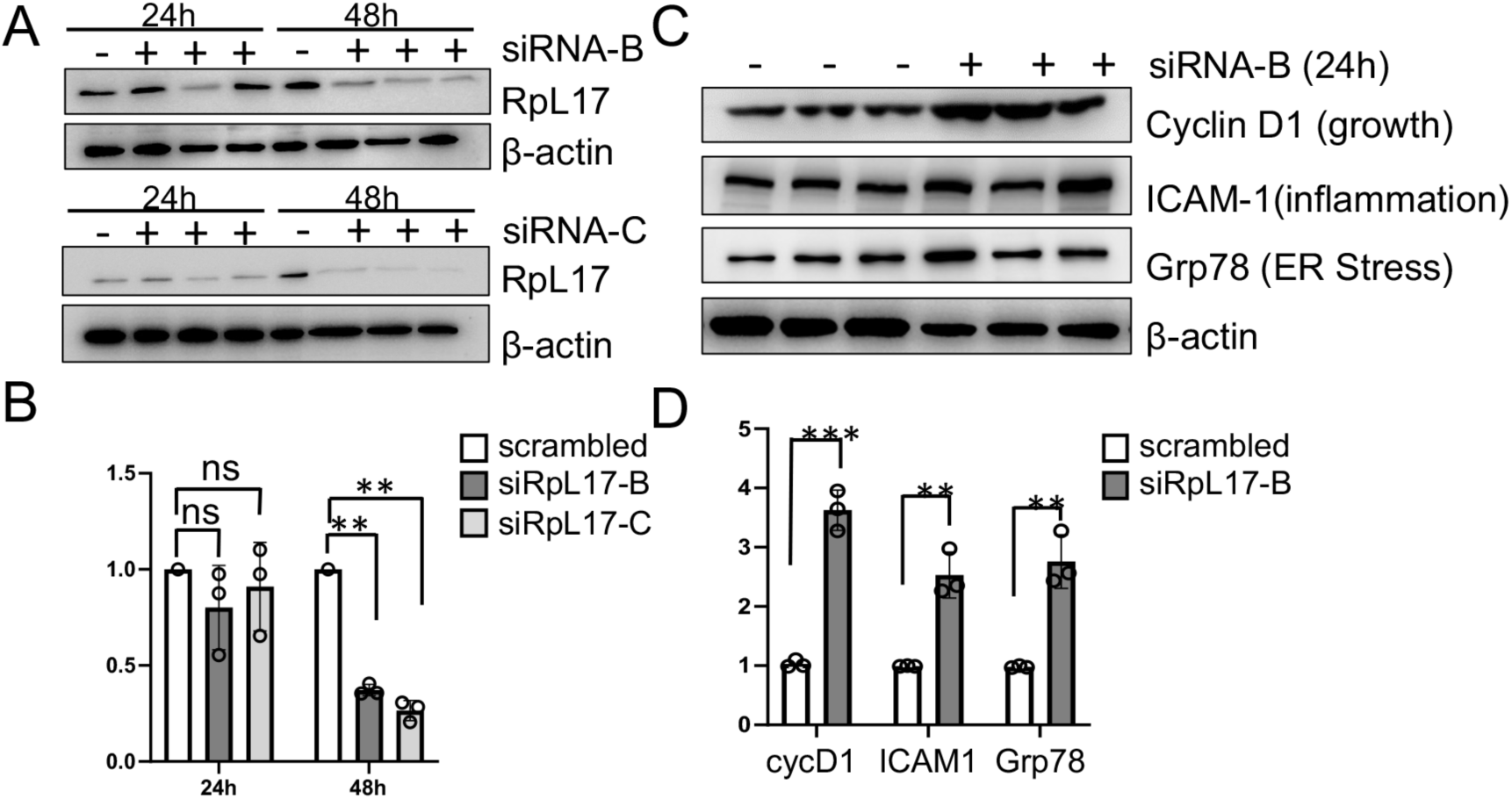
Human EC with low RPL17 have similar stress response to mouse RpL17-Het EC. (A) RpL17 was depleted from human aortic EC (HAEC) using siRNA. Western blots of lysates were probed as indicated and quantified in (B). (C) HAEC depleted of RPL17 were probed for cyclin D1, ICAM-1, and Grp78; all normalized to beta-actin (quantified in (D)).

## Discussion

The major finding of this study is the discovery of a novel mechanism for the formation of intima-media thickening caused by decreased expression of the ribosomal protein RpL17, which we have demonstrated both genetically and in response to disturbed flow (d-flow, Fig. 1 and Fig. S1). Mechanistically, we found that decreased RpL17 increased ER stress and activated the integrated stress response (ISR), which was associated with a global decrease in translation, but increased translational efficiency of proteins involved in redox, ER/UPR stress, glycolysis, matrix and migration, inflammation, and EndMT. These findings are novel because most studies investigating the effects of flow on EC focus on transcriptional regulatory changes, not translational. Furthermore, decreases in EC global translation were associated with altered translational control: specifically increased translation initiation of cap-independent mRNAs. In the presence of d-flow and changes in translation, EC were “stimulated” leading to proliferation, caused in part by increased expression of the Mdm2 ligase that ubiquitinates the growth suppressor p53, increasing its proteosomal degradation; thereby eliminating a major barrier to proliferation.

The present data provide a mechanistic explanation for the recent GWAS study that identified RpL17 as a locus of increased risk for metabolic syndrome (waist-to-hip ratio), which is a key factor in CAD ^32^. To our knowledge, only one other ribosomal protein, MRPL14, has been associated with CAD ^33^. In summary, these results indicate a key role for RpL17 in translational control that maintains normal EC phenotype and function. Furthermore, our data provide a novel mechanism for IMT: D-flow decreases RpL17 expression that inhibits EC protein translation due to ER stress and the integrated stress response (ISR), which leads to metabolic reprogramming, EndMT, and IMT (Fig. 8) ^34,35^. A key finding was the presence in RpL17-Het EC of changes in expression of proteins associated with endothelial dysfunction. These included increased expression of proteins that cause oxidative stress, inflammation, migration, EC barrier disruption, EndMT, and metabolic reprogramming to a glycolytic state; as well as decreased expression of inhibitors of proliferation and proteins necessary for oxidative phosphorylation.

**Figure 8:**
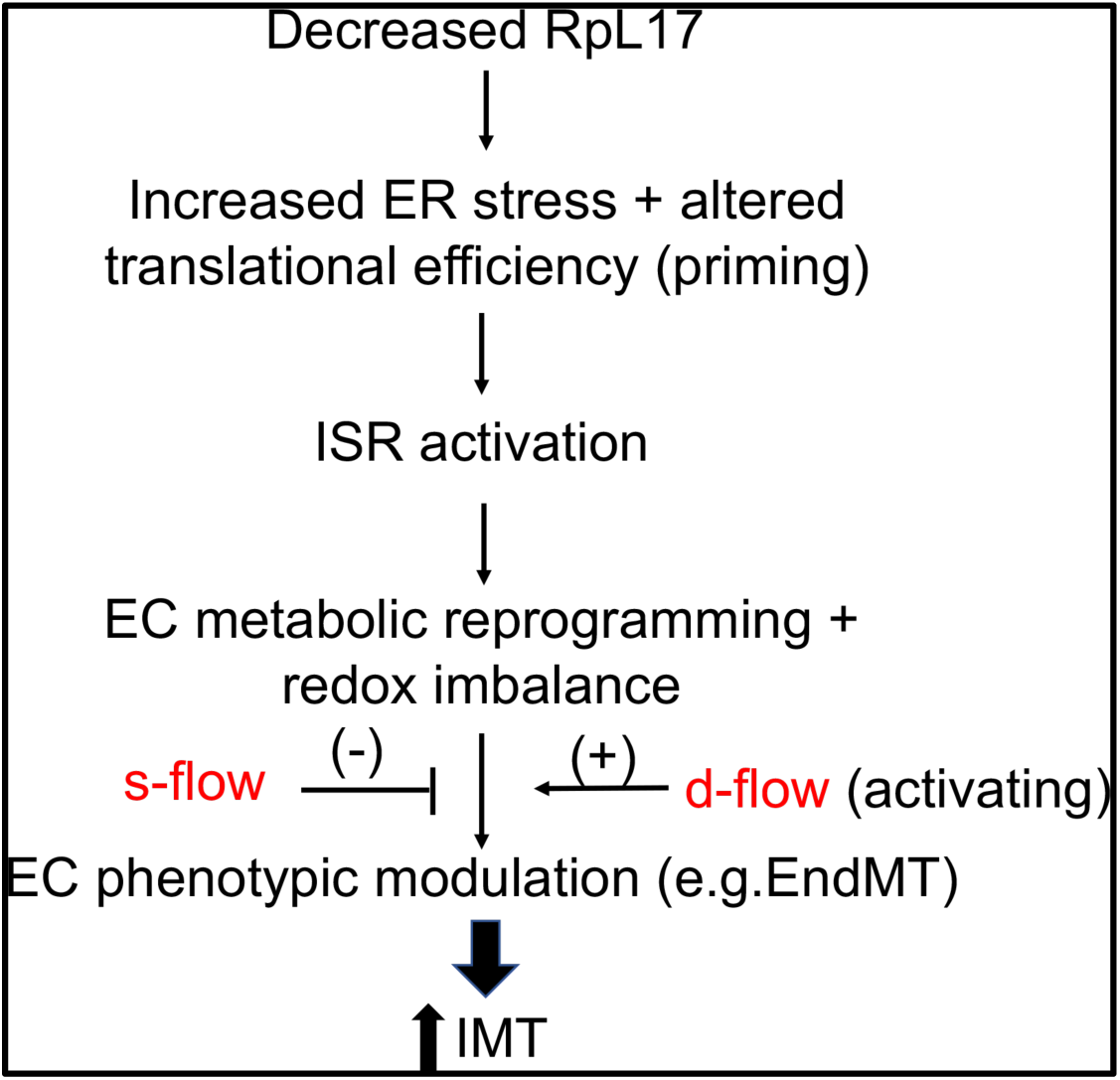
Model diagram of proposed molecular mechanism for RpL17-mediated translational control of endothelial cells exposed to laminar flow. When levels of RpL17 are decreased in EC, translational control shifts as identified by differential polysome-associated mRNAs in EC-RpL17. These changes activate ER stress and the stress-responsive eIF2 kinase, PERK. ER stress has been shown to induce EndMT. Consistent with PERK activation, levels of phospho-eIF2 alpha and the ISR-dependent transcription factor Atf4 increase in RpL17+/- EC, indicating ISR activation. Metabolic and proteostatic changes in EC-RpL17 cells correlate with changes observed in translational efficiency of key regulatory enzymes in the same pathways. Combined, these alterations lead to pathological IMT and increased susceptibility to atherosclerosis.

### Translational efficiency changes in RpL17-Het EC

Translational efficiency is a rapid and energy-efficient mechanism to respond to environmental changes that occur frequently to EC. Importantly, it decreases the need for mRNA synthesis, which requires much more energy and time. Changes in translational efficiency are usually caused by altered translation initiation (cap-independent), and by interactions of mRNA with microRNAs, RNA binding proteins, and long non-coding RNAs ^36,37^. The mRNAs chosen for measurement of translation efficiency showed significant cap-independent regulation of translation. Common mechanisms involve recruitment of the pre-initiation complex by internal ribosome entry site (IRES) elements and their IRES transacting factors and increased initiation of translation from 5’ upstream open reading frames (uORFs). We investigated the human IRES database that has curated experimental validation for IRES elements and uORFs. The sequences were significantly enriched in the RpL17-dependent translational efficiency target mRNAs. These results demonstrate that eIF2 alpha-independent translational control is an essential feature of the EC response to stress, such as occurs with d-flow.

### D-flow stimulates ROS and activation of the ISR in RpL17+/- EC

The cardiovascular system is especially responsive to biomechanical forces compared to other systems ^38^. Existing data for the EC response to changes in shear stress (e.g. d-flow vs. s-flow) have been shown to alter physiologic processes such as vasodilation, inflammation, permeability, and cell proliferation ^26,39,40^. This study shows significant modification of the EC response to d-flow by cap-independent translational control ^38,41^. While it is very likely that smooth muscle cells and inflammatory cells play an important role in vascular remodeling, the present study shows a cell autonomous role for EC in responding to d-flow. Increased ROS is probably the most common cause of EC phenotypic change due to its multiple effects on key physiologic functions. In this study, decreased RpL17 increased ROS via several mechanisms: 1) Increased ER stress and the UPR; 2) switching to glycolysis because of a shortage in key proteins necessary for mitochondrial complexes I and III; and 3) redox imbalance due to increased translation efficiency of prooxidant enzymes such as Nox4 (Fig. 6)^29,42^.

### EndMT is enhanced by combined actions of d-flow and low RpL17 expression

EndMT and inflammation are fundamental aspects of pathologic vascular remodeling, especially when induced by d-flow ^11,22,43^. Here we show for the first time that chronic ISR and its metabolic changes “prime” EC for EndMT by significant inhibition of the steady-state transcriptome and proteome, which maintains differentiated EC function. These primed EC are then “stimulated” by d-flow via genetic and epigenetic pathways associated with migration, proliferation, and inflammation ^44^. Two key pieces of data from RpL17-Het EC strongly support a pivotal role for EndMT to promote the vascular remodeling induced by low RpL17 combined with d-flow. First, we found that Snail, a transcriptional repressor of E-cadherin, and Gata4, a transcriptional activator of EndMT genes, showed increased translational efficiency in RpL17-Het EC. Second, increased immunostaining for both Snail and pSmad2/3 that are downstream targets of the EndMT-promoting TGF-beta signaling pathway, were observed in RpL17-Het PCL carotids (Fig. 6). Support for this concept comes from the single-cell RNA-seq data of the Jo lab in which EC exposed to d-flow after PCL showed features of EndMT, with decreased Klf2, and increased inflammation and glycolysis ^11,25^. Our data extend these findings because we have discovered that IMT induced by d-flow requires a chronic translational, ISR-dependent program, in addition to the well-described epigenetic and genetic programs ^44^ (Fig.2, Fig.8).

### Increased EC proliferation in Rpl17 heterozygotes mediated by ISR-dependent translation

There are several potential mechanisms for EC proliferation with decreased ribosome number and translation: 1) selective cap-independent, IRES-dependent translation of growth-pathway proteins such as cyclin D1 and PCNA; and 2) increased expression of Mdm2 that ubiquitinates the growth suppressor p53, increasing its proteosomal degradation. Metabolic shifts in ribosomopathies (usually mutations in ribosomal proteins) show similar changes to RpL17-Het EC, especially increased glycolysis, redox stress and proliferation ^45,46^. Increased glycolysis is closely linked to proliferation, as is the case in EC and cancer cells ^47–51,52^. Further support for this idea is that ribosomopathies such as Diamond Blackfan anemia and Treacher-Collins syndrome have been linked to increased cancer risk ^53,54^. Furthermore, ribosomal protein genes are frequently mutated in cancer, occurring in 43% of the most common cancers ^55^. RPL17 is among the top ten most mutated ribosomal proteins in human cancers (found in ∼19% of cancers). What is unique about RPL17 that that causes it to be associated with so many diseases that involve cell growth? The answer will require much further investigation. But some potential causes are 1) Its presence in the exit tunnel, which may cause more protein misfolding and more ER stress; 2) An unidentified extra ribosomal function, since it has unique subcellular locations in both EC and smooth muscle cells compared to other ribosome subunit proteins; 3) Its requirement for ribosome maturation, which leads to decreased ribosome numbers and polysomes, causing perhaps more severe ISR than other ribosomal proteins; and 4) its expression is flow-responsive.

In summary, our data show that translation is an important regulator of EC phenotype: EndMT, EC proliferation and inflammation, and fibrosis all cause IMT (Fig. 8). The most important mechanism altered in RpL17-Het EC appears to be activation of the ISR. It changes translational efficiency, which is a rapid and low-energy process that can be easily and tightly regulated by cells under stress. Our data point to regulators of translational efficiency as novel candidates to treat cardiovascular diseases such as carotid IMT. Potential regulators include stimulus (d-flow vs s-flow), cell (EC vs. smooth muscle cells, inflammatory cells), RNA binding factors (proteins, microRNAs, long noncoding RNAs) and proteins (Nox4 and Mdm2).

## Acknowledgements

The authors gratefully acknowledge the expert assistance with electron microscopy from Drs. Karen Bentley and Chad Galloway, and Ms. Kelsea Cristillo; with quantification of RpL17 in mouse carotids from Mr. Sean Hopkins; with sectioning from Erika Flores Medina; and from Ms. Sharon Senchanthisai and Ms. Amanda Pereira with mouse genotyping, maintenance, and tissue collections and preparation.

## Supplemental Material S1: Methods

### Mice

Mice were maintained under a standard 12-hour light-dark cycle with regular chow and water *ad libitum* in accordance with institutional UCAR standards. The floxed RpL17 mice were generated by creating founder chimeras with targeted heterozygous p5 male JM8 C57BL/6N embryonic stem cells (ESCs) purchased from the European Mouse Mutant Cell Repository (EuMMCR) as part of EUCOMM (MGI Allele ID# 2448270, Clone IDs HEPD0801_5_E04 and HEPD0801_5_H03, allele name *Rpl17^tm3a(EUCOMM)Hmgu^*; ^56^). ESCs were expanded and injected into blastocysts by the URMC Transgenic Facility to generate founder mice. Floxed RpL17 heterozygotes were maintained on a C57 BL/6 background, and bred to Cdh5-Cre (stock #017968, strain B6;129-Tg(Cdh5-cre)1Spe/J from Jackson Labs) transgenic mice ^57^.

### Translational efficiency analysis

Translational efficiency analysis was performed on sets of 21 fractions collected from both RpL17-WT and RpL17+/- MLMEC. Fractions were pooled as follows: Fractions 1 & 2, ribosome-free; fractions 3 & 4, 40S & 60S subunits; fractions 5 & 6, 80S monomers; fractions 7 & 8, 80S dimers; fractions 9-15, light polysomes; fractions 16-21, heavy polysomes. An equal volume (375 µL) of each ribosomal fraction was used to extract mRNA using Trizol LS (Invitrogen) according to the manufacturer’s recommended protocol. Prior to the RNA extraction, 1 ng of *in vitro* transcribed Renilla luciferase mRNA was spiked into each fraction as a normalization Control for RNA content. An equal volume (7.5 µL) of purified total RNA (range: 90-430 ng/µL) was reverse transcribed using iScript cDNA synthesis (Bio-Rad). Quantitative real-time PCR amplification of transcripts was performed using 4 µL diluted cDNA (range: 13-64 ng RNA equivalents of cDNA), 1 µM specific primers (see supplemental methods), and SybrGreen PCR reagent (Bio-Rad). Cycle parameters were a 2-step protocol of 95°C denaturation and 60°C annealing/extension for 40 cycles. Luciferase amplification served as an internal Control.

### Statistical analysis

Statistical analysis was performed using GraphPad Prism 9. Quantitative data was collected by an experimenter blinded to the genotype or treatment of sample. For fluorescent image analysis, Fiji/ImageJ2 (v.2.3.0) image processing software (NIH) was used on single-channel images with the same threshold applied per experiment, and mean pixel density was recorded per image. Unpaired 2-tailed Student *t*-test with Welch’s correction was used when comparing two groups; with more than two groups, 2-way ANOVA with Tukey’s multiple comparisons test was used. A value of *p* < 0.05 was considered statistically significant. Replicates are biological and indicate number of mice or cultures used, except for subcellular structures (e.g. ribosome particle density), in which case the number of fields examined is equal to *n.* Error bars on graphs indicate the standard deviation of the sample group.

### Vessel morphometry

To determine the volume of vascular compartments, carotid arteries were paraffin embedded, and serial sections collected every 200 microns from the bifurcation point toward the aorta. Sections were stained with hematoxylin & eosin, and light micrographs collected on an Olympus BX41 equipped with a Spot digital camera. Images were analyzed using Fiji (NIH), and volumes of each compartment quantitated. Stenosis was calculated as (I + M)/EEL as described.^1^

### Antibodies and reagents

The following antibodies were used for immunostaining and western blotting: RpL17 (Thermo Fisher #PA529397), CD144 (BD Biosciences #555289), BiP (BD Biosciences #610978), Chop (Thermo Fisher #MA1-250), p-eIF2a (Cell Signaling Tech #3398), pPERK (Invitrogen #MA5-15033), Pfkfb3 (Proteintech #13763-1-AP), 8-oxo-dG (Trevigen #4354-MC-050), 4HNE (EMD Millipore #393207), PCNA (Dako PC10, #M0879), ICAM (ThermoFisher YN1/1.7.4 #14-0541-85), VCAM (GeneTex #GTX14360), CD45 (BD Bioscience #550539), CD31 (Abcam #ab28364), Nox4 (Abcam #ab154244), Activating transcription factor-4 (Atf4; Abcam #ab85049).

### Immunofluorescence

For immunofluorescence of mouse carotid sections, sections were deparaffinized, antigen retrieval performed using citrate buffer pH 6.0 (EMD Millipore), and blocked with 5% normal goat serum (NGS), 0.1% Triton X-100 in phosphate-buffered saline (PBS) at room temperature. Primary antibodies were diluted in antibody diluent buffer (Dako) and incubated overnight at 4°C. Alexa Fluor fluorescent conjugated secondary antibodies (goat anti-rat/rabbit/mouse 488/546/647) were diluted 1:500 in 5% NGS/PBS, and sections were mounted in aqueous mounting medium containing DAPI (DAPI-Fluoromount G, Southern Biotech) and coverslipped. Confocal images were collected on a laser-scanning confocal microscope (Olympus IX51) equipped with Olympus FluoView imaging software (FV1000).

### MLMEC culture

Mouse lung microvascular endothelial cells were harvested from mice under ketamine/xylazine anesthesia. Briefly, mice were perfused intracardially with 0.9% saline, and lungs were inflated with DMEM containing 2 mg/mL collagenase. Lungs were removed and incubated in a 37C waterbath in 2 mg/mL collagenase solution for 30-40 minutes, with vortexing every 10 minutes. DMEM + 20% FBS was added 1:1 to dissociated lung solution, and cells filtered through a 70-micron mesh. Lung EC were purified by magnetic positive selection using anti-CD31-conjugated beads (Invitrogen sheep anti-rat IgG Dynabeads, BD Biosciences rat anti-ms #555289), and plated on gelatin-coated dishes in DMEM containing 20% serum.

### Cell cycle analysis

Cultured Control and floxed RpL17 MLMEC were infected with lentiviral Cre-EGFP, and DNA stained with DRAQ5 (647ex/681em) for nuclear DNA content/cell cycle phase analysis. Cells were analyzed by flow cytometry, gated based on GFP fluorescence and analyzed for percentage of cells in G0/G1, S, or G2/M phase relative to the total GFP positive cells.

### Western blots

Cultured EC were scraped into ice-cold PBS, pelleted, and lysates prepared using Cell Lysis buffer (Cell Signaling Technology) containing phosphatase and protease inhibitors (Millipore-Sigma). Lysates were resolved through Tris-glycine gels and blotted to nitrocellulose, blocked with 5% dried milk, and reacted overnight with primary antibodies diluted in 2% BSA/PBS-Tween (0.1%). HRP-conjugated secondary antibodies (Jackson Immunoresearch) were reacted at room temperature for 1-2 hours and blots were developed using chemiluminescence substrate (EMD-Millipore) and imaged using a BioRad Gel-Doc imaging system.

### Partial carotid ligations

Carotids of male and female mice were ligated between 10-12 weeks of age as previously described^4^. Briefly, mice were anesthesized with isofluorane and maintained on a heating pad at 37°C. The left carotid artery was exposed, and both the external and internal (superior to the occipital artery) carotid arteries were ligated with silk suture thread, leaving the occipital artery patent. Control sham surgeries were performed in an identical manner, placing suture under the carotids but not ligating the thread. Mice were sacrificed 14 days post-ligation, perfused with 10% neutral-buffered formalin, and both left and right (control) carotids were collected for immunostaining and histology.

### Polysome profiling

Ribosomes were purified from MLMEC. A 15-cm dish of 90-100% confluent cells was incubated in in culture medium containing 100 µg/mL cycloheximide for 15 min at 37°C. Cells were scraped into ice-cold PBS + cycloheximide, and pelleted. Cells were resuspended in TMK lysis buffer (10 mM Tris-HCl 7.4, 5 mM MgCl_2_, 100 mM KCl, 1% Triton X-100, 0.5% deoxycholate, 1 U/mL RNase inhibitor, 2 mM DTT, and 0.1 mg/mL cycloheximide), incubated on ice 20 min, and centrifuged at 16,000 x g at 4°C for 15 minutes. The protein concentration of supernatants was measured using the Bradford assay (BioRad) and equivalent amounts loaded onto 10-50% continuous sucrose gradients. Lysates were resolved at 50,000 x g for 4 hours, and 1 mL fractions collected on an ISCO density gradient fractionator.

### Oxidative stress, HNE, 8-oxo-dG and SOD2

All measurements for oxidative stress were performed by immunostaining of carotid or aortic sections. For SOD2, *en face* immunostaining of EC was performed on the aortic arch.

### Metabolism analysis

The Seahorse XFe96 extracellular flux analyzer (Agilent) was used to measure mitochondrial and glycolytic activity of MLMEC. Cells were seeded at a density of 3 x 10^4^ cells per well in an XF96 cell culture microplate in DMEM media and incubated overnight. Next day, the media was discarded, cells were washed with PBS and incubated in Seahorse XF base medium (Agilent, 103334-100) (Santa Clara, CA, USA) containing 2g/L glucose, 1X glutamax (Thermo Fisher Scientific, 1798324) and 1X sodium pyruvate (Thermo Fisher Scientific, 11360070) for 3 hours in CO2 free incubator. Oxygen consumption rate (OCR) and extracellular acidification rate (ECAR) were measured following with sequential injection of 2 uM oligomycin, followed by 1 uM FCCP, and a final injection of rotenone and antimycin A (0.5 uM). Oligomycin (O4876), carbonyl cyanide-4-(trifluoromethoxy)phenylhydrazone (FCCP, C2920), rotenone (R8875), antimycin (A8674), and 2-deoxy-D-glucose (D6134-5G) were purchased from Sigma-Aldrich (St. Louis, MO).

### Lactate measurement

MLMECs were seeded in six-well plates (3 × 10^5^ cells/well) and lactate dehydrogenase (LDH) in the supernatant was measured using L-Lactate Assay Kit (#1200011002, Eton Bioscience, San Diego, CA, USA) according to the manufacturer’s protocol.

### Aortic en face preparations

Mouse aortas were collected following ketamine/xylazine anesthesia, intracardiac perfusion with heparinized saline then neutral-buffered formalin (10%), dissection and overnight incubation at 4C in blocking buffer (1% bovine serum albumin (BSA) + 1% Triton X-100 in PBS). Primary antibodies (RpL17: Thermo Fisher #PA529397; CD144: BD Biosciences #555289) were diluted 1:200 and 1:250, respectively, in 0.1% BSA in PBS, and aortas were incubated with primary antibodies overnight at 4C with gentle agitation on a rocking platform in horizontal Eppendorf tubes. Aortas were rinsed three times with PBS + 0.1% TritonX-100 with agitation, and incubated 1 hr at room temperature in Alexa Fluor-conjugated secondary antibodies diluted 1:500 in 3% NGS + 0.1% BSA in PBS. Post-secondary antibody washes were performed with PBS and mounted in ProLong Gold anti-fade mounting media.

Laser-scanning confocal imaging was performed on an Olympus IX51 microscope equipped with Olympus FluoView imaging software (FV1000); all images were collected as z-stacks delimited by the CD144+ endothelial layer, then summed into a maximum projection image.

### HUVEC and HAEC Cell Cuture and siRNA transfection

ECs were isolated from human umbilical veins (HUVEC) as described (RRR). Cells were grown in Medium 200 (Thermo Fisher Scientific) supplemented with 5% FBS (HyClone), heparin (Sigma), and LSGS (S-003-10). Cells were treated Lipofectamine RNAiMax (Invitrogen,13778-075) and 50nM siRNAs in OptiMEM (Gibco, 31985-070). Human siRpL17 (Silencer Select siRNA vs Rpl17. ID s54963 and control negative siRNA (4390843) were purchased from Thermo Fisher Scientific. Lifeline® normal Human Aortic Endothelial Cells (HAEC) (FC-0014) were grown in Lifeline® VascuLife® Medium.

### Electron microscopy

For electron microscopy, Control and Rpl17+/- MLMEC were plated at 3 x 10^5^ cells/dish in 6 cm dishes containing 18 mm glass coverslips coated with 2% gelatin/PBS. Cells were incubated for 48 hr at 37C to confluency, then media removed and replaced with room temperature fixative containing 2.5% glutaraldehyde + 4% paraformaldehyde at room temperature for 1 hr. Coverslips were then placed at 4C in fixative and further processed in osmium tetroxide and imaged at 60,000x/0.2 um resolution with a transmission electron microscope in the URMC Electron Microscopy Core Shared Resource Laboratory. Quantification of endoplasmic reticulum area and sac width were performed with EM images in Fiji (Image J).

### Puromycylation assays

Prior to labeling actively-translating proteins on ribosomes with puromycin, HUVEC cells on 2% gelatin-coated dishes (+/- coverslips) at 70% confluency were transfected with scrambled or siRNA directed against human RPL17 (100 ug siRNA per 10 cm dish) in Lipofectamine RNAi Max reagent for 6 hr, then transfection media was removed, and cells grown to confluency. For puromycylation labeling, methods were used as reported previously ^58,59^ with slight modifications. At the time of labeling, 91 uM puromycin + 208 uM emetine was added to HUVEC media, and cells were kept 15 minutes at 37C. Media was removed, and HUVEC were placed on ice. For confocal imaging, cells on coverslips were treated with permeabilization buffer (50 mM Tris-HCl pH 7.5, 5 mM MgCl_2_, 25 mM KCl, 355 uM cycloheximide + protease inhibitors) for 2 min, washed with 50 mM Tris-HCl pH 7.5, 5 mM MgCl_2_, 25 mM KCl, and 0.2M sucrose, then fixed with 3% paraformaldehyde in PBS for 15 min at room temperature. Cells were rinsed with PBS, blocked in buffer containing 0.05% saponin, 10 mM glycine, 5% NGS in PBS for 15 minutes, then incubated with anti-puromycin (ms Sigma #MABE343, 1:500) + anti-RPL17 (rb Thermo Fisher #PA529397, 1:250) diluted in blocking buffer for 1 hr at room temperature. Alexa Fluor-conjugated antibodies were diluted 1:500 in 3% NGS/PBS, and coverslips were mounted in ProLong Gold anti-fade mounting medium for confocal imaging.

For western blotting, cells were harvested in ice-cold lysis buffer (Cell Signaling Tech #9803) containing protease inhibitors, sonicated briefly, and cell debris removed by centrifugation. 25 ug of the puromycylated HUVEC lysates were separated on 10% SDS-PAGE gels and blotted to nitrocellulose; blots were then blocked with 5% milk/PBS/0.1% Tween-20 and probed with anti-puromycin antibody #MABE343 diluted 1:10,000 in 2% BSA/PBS.

### RNA sequencing

For total RNA sequencing, Control and Rpl17+/- MLMEC (P3) were grown to confluency in 10 cm dishes, rinsed with D-PBS, then placed on ice and scraped into ice-cold D-PBS. Cells were pelleted, then resuspended in 350 uL QIAGEN RLT buffer + DTT, placed on dry ice, and processed for sequencing by the URMC Genomic Research Core facility.

### IRES, uORF, and rG4 5’ UTR analysis of mRNAs

RpL17-dependent translational efficiency target mRNAs were analyzed for IRES and uORFs using the Human IRES Atlas database search engine (http://cobishss0.im.nuk.edu.tw/Human_IRES_Atlas/QueryGene) ^60^ for putative and experimentally-validated IRES elements and upstream ORFs; and the rG4 database search engine QRGS Mapper (https://bioinformatics.ramapo.edu/QGRS) ^61^ for G4 quadruplexes found in RNAs.

## Supplemental Material S2: Figures

**Figure S1:**
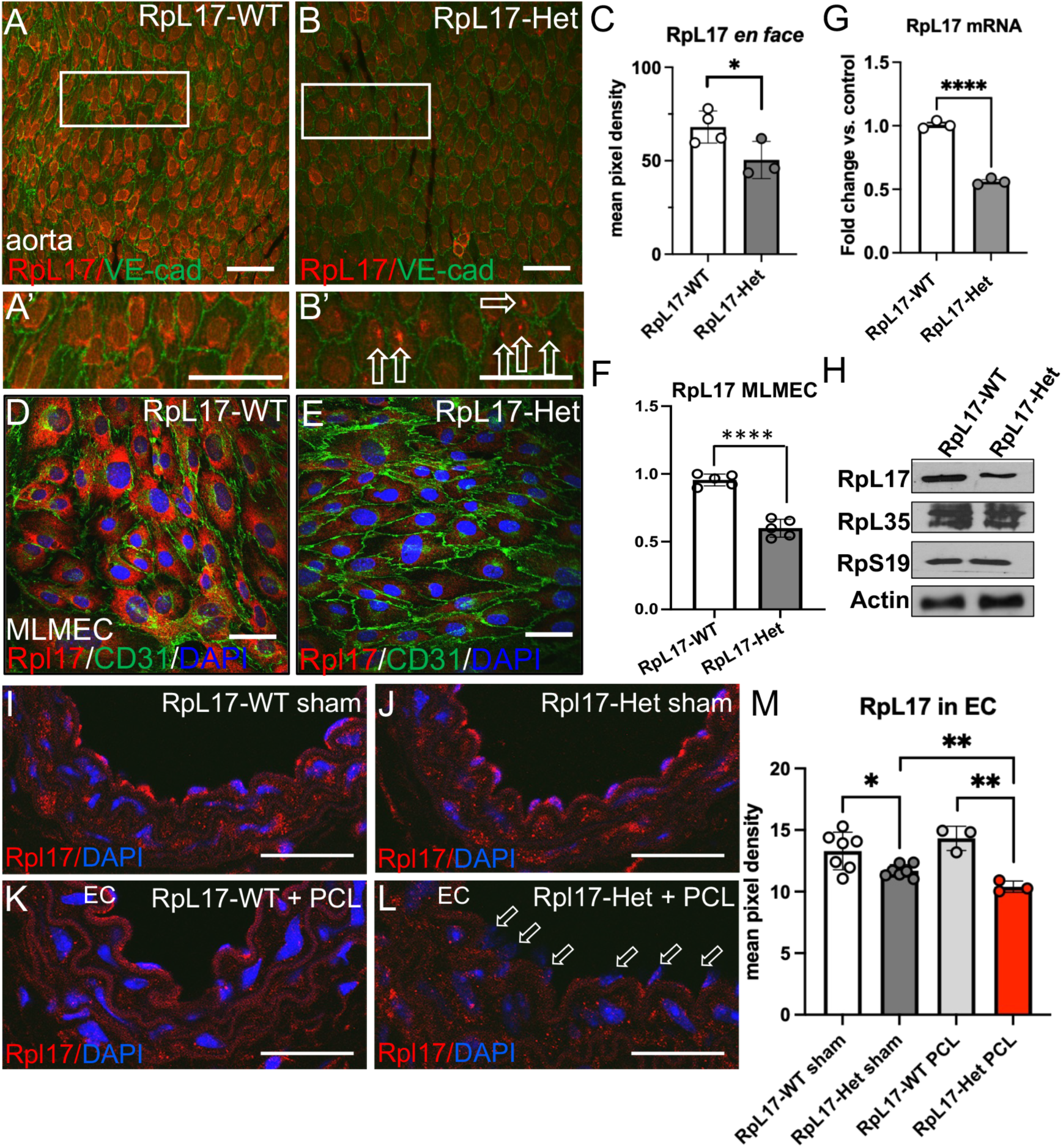
RpL17 heterozygous EC have reduced RpL17 mRNA and protein. (A) RpL17 mRNA in EC-RpL17 MLMEC measured by qPCR. (B-D) Immunostaining of cultured MLMEC with anti-RpL17 (quantified in D). (E) Western blot of ribosomes from MLMEC show reduced RpL17 but no change in RpL35 and RpS19 levels (n=3 separate lung isolations from 3 pooled mice of each genotype; p<0.05). (F-H) Decreased RpL17 expression in EC-RpL17 aortic EC. *En face* immunostaining of RpL17 in descending aorta of EC-RpL17 (G) vs. control aorta (F); quantified in H (n=4 controls/3 hets; p< 0.01). Scalebars are 50 microns.(I-M) RpL17 immunostaining in mouse carotid sections for RpL17 identified strong positive staining in RpL17-WT EC under steady flow (I) and in RpL17-WT after PCL (d-flow) (K). (J) RpL17-Het PCL carotid before PCL identified a decrease in RpL17 which is decreased further in the presence of d-flow in RpL17-Het carotids following PCL (L). (M) Quantification of RpL17-immunostained carotid sections shown in A-D. *\*, p< 0.05; **, p< 0.01. N=7 (sham), 3 (PCL)*.

**Fig. S2:**
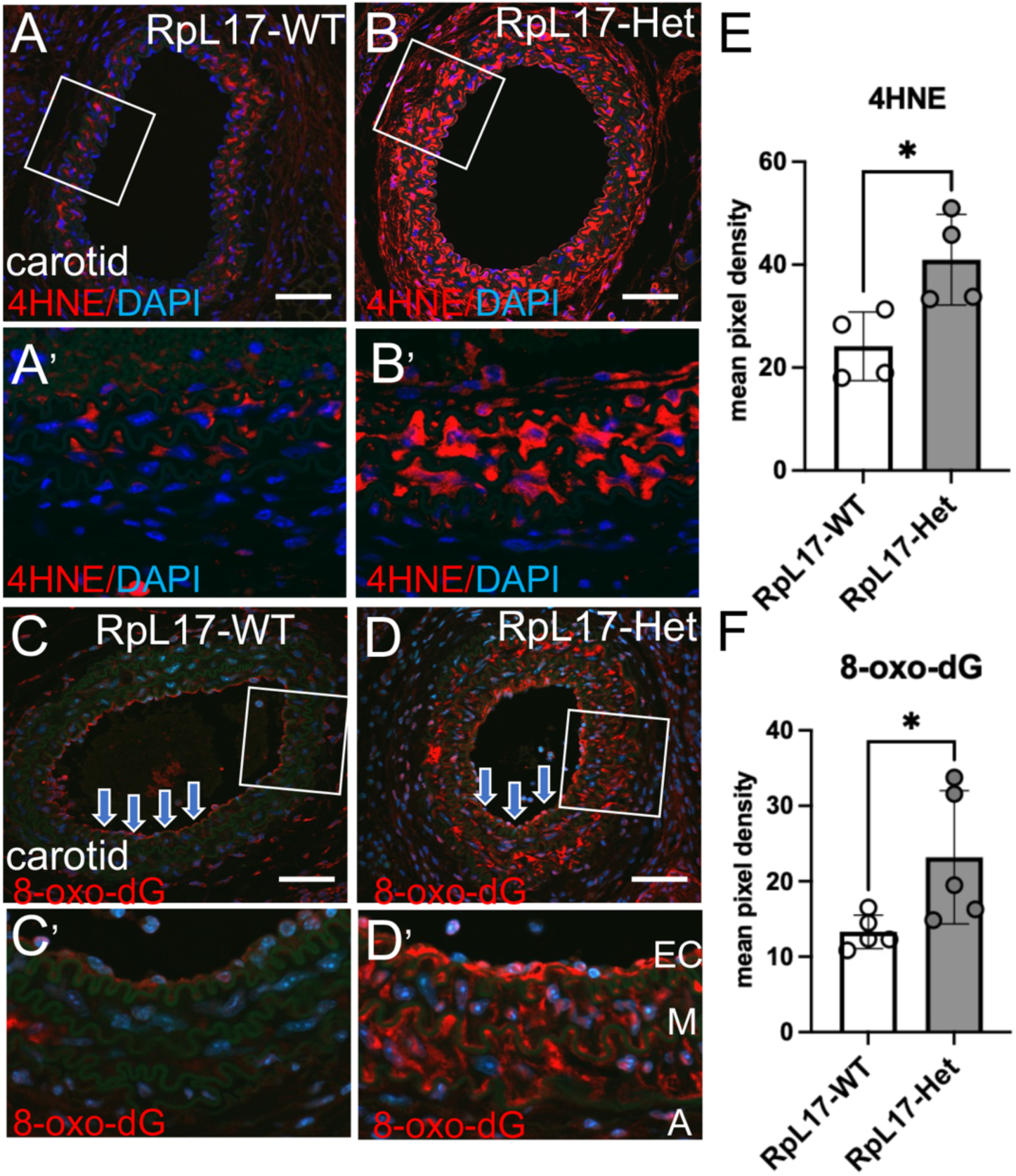
Oxidative stress is increased in EC-RpL17 exposed to d-flow. (A,B) Immunostaining of sections from ligated carotids with anti-4-hydroxynonenal (4HNE), a marker of lipid peroxidation, in RpL17-Het vs. RpL17-WT (compare B vs. A, quantified in E). (C,D) Immunostaining of carotid sections with anti-8-hydroxydeoxyguanosine (8-oxo-dG) in RpL17-Het vs. RpL17-WT (D vs. C, quantified in F). Scalebars are 50 microns.

**Figure S3:**
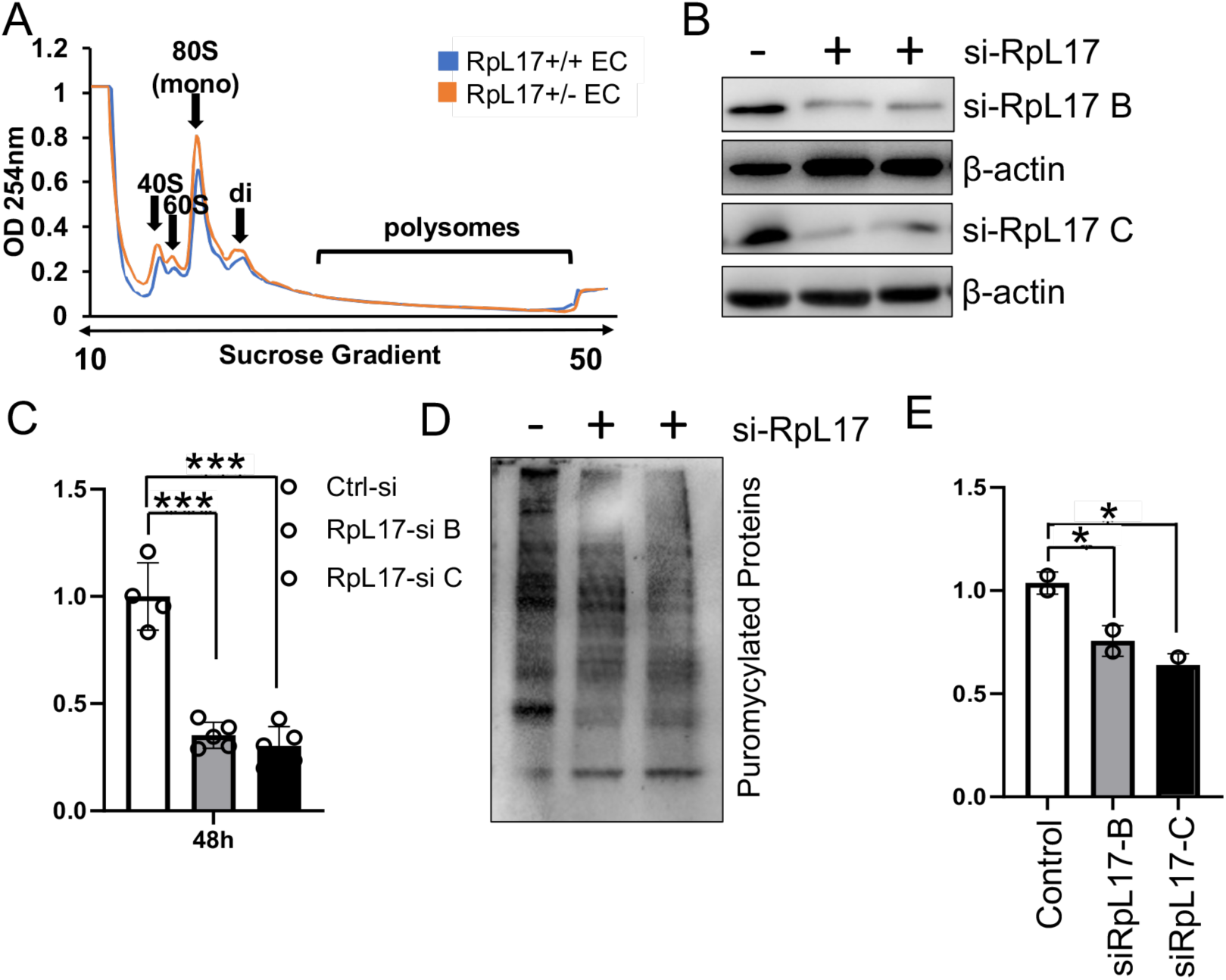
Altered polysome profile and a reduction in global translation with reduced RpL17 in EC. (A) Polysome profiling of MLMEC from RpL17-WT vs. RpL17-Het. Note the slight increase in free 40S and 60S subunits, monosomes, and disomes, consistent with cell stress and overall reduction in translation. (B) Western blot of lysates from HUVEC +/- siRpL17 (two different siRNA against RpL17, RpL17-siB or -siC), labeled with puromycin, and probed for Rpl17 showed decreased RPL17 protein levels with siRpL17 (quantitated in C). (D) Western blot of HUVEC lysates following puromycylation probed with anti-puromycin Ab (quantified in (E); p<0.05). Reduction in siRPL17 HUVEC indicates reduced global translation.

**Figure S4:**
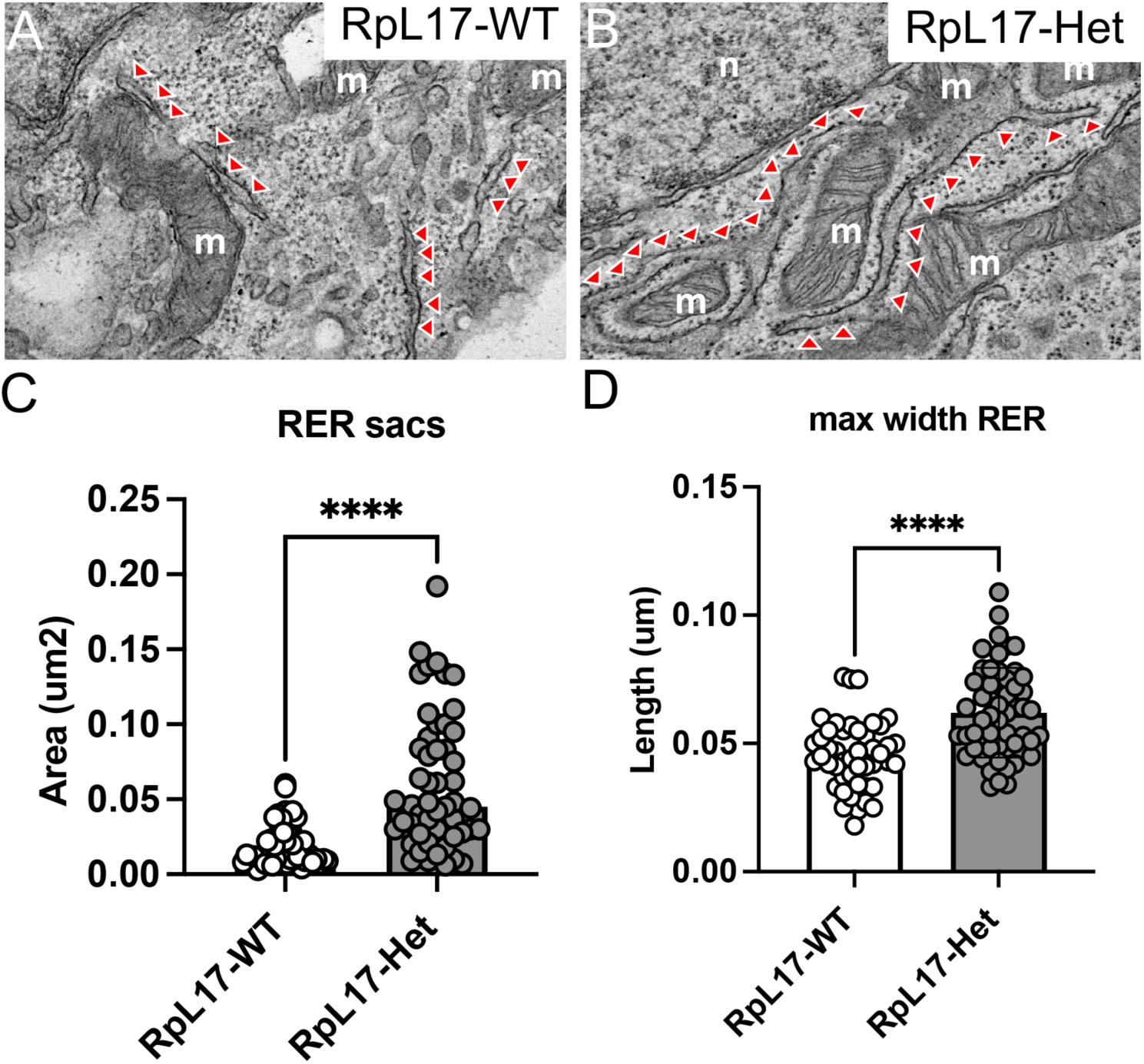
RpL17-Het MLMEC rough endoplasmic reticulum (RER) sacs are increased in size. (A-B) EM micrographs of cultured MLMEC containing rough endoplasmic reticulum (RER, red arrowheads). Quantification of EM images revealed increased maximal width of RpL17-Het RER sacs vs. RpL17-WT (D), and increased area of RpL17-Het RER sacs (C). No significant difference was found in the number of caveolae near the plasma membrane (data not shown). *N=3 samples per genotype; each dot represents one sac. ****, p< 0.0001*.

**Figure S5:**
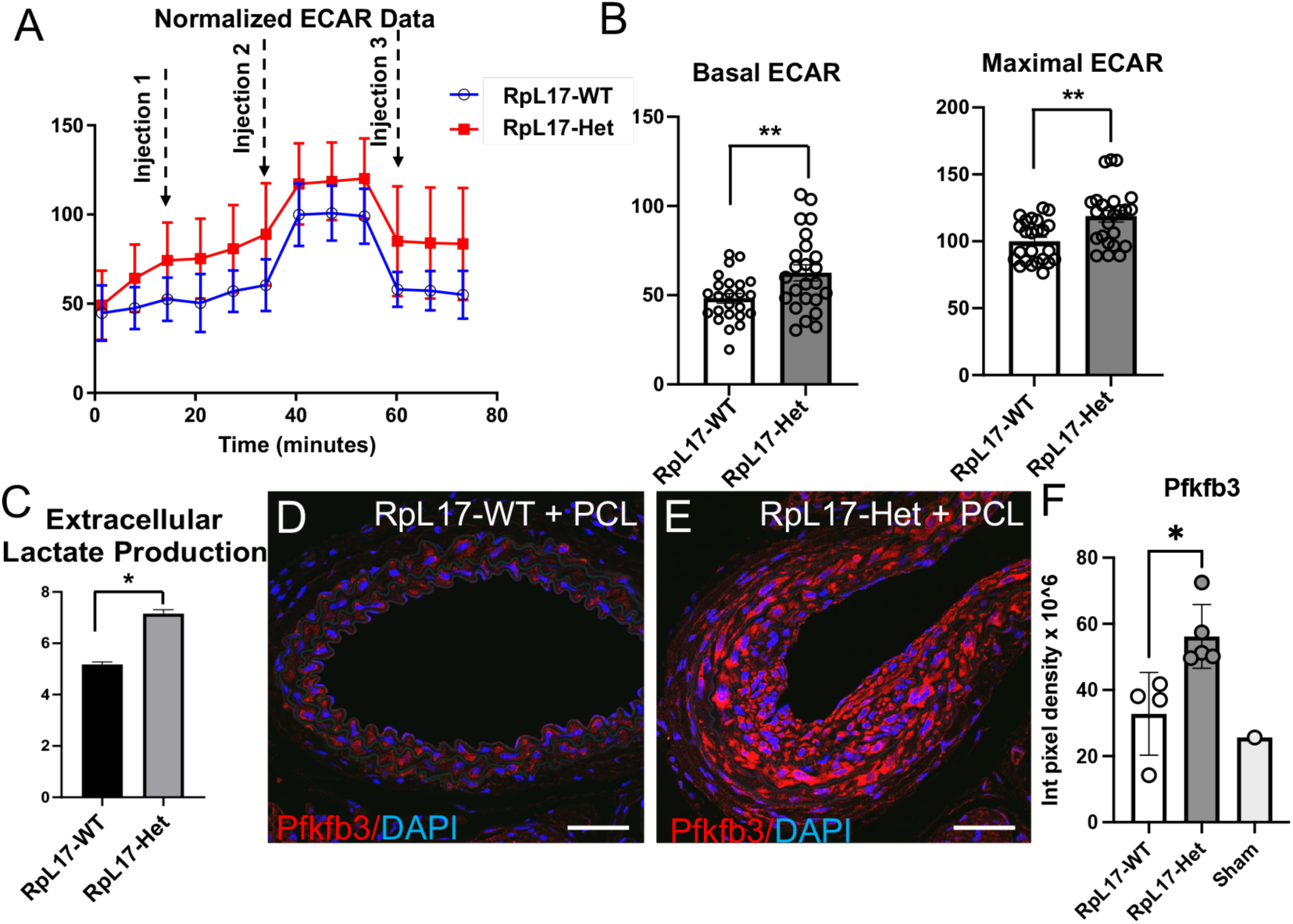
Decreased RpL17 levels in RpL17+/- EC favors glycolysis. (A) RpL17-Het MLMEC increased extracellular acidification rate or ECAR (≈ 26%) as compared to RpL17-WT MLMEC. Seahorse analysis was used to measure ECAR; injection 1: 1µM oligomycin; injection 2: 2µM FCCP; injection 3: 1µM (rotenone + antimycin A); FCCP; carbonyl cyanide p-trifluoro methoxyphenylhydrazone. (B) Indicates increased ECAR at both basal and maximal (B) states in RpL17-Het. (C) Increased extracellular lactate production in RpL17-Het MLMEC compared to control. Data is indicative of increased glycolysis in RpL17-Het (Average of n=8 for A, B; n=2 for (C; p< 0.05). (D,E) Anti-Pfkfb3 (glycolytic enzyme) immunostaining of carotid sections post-ligation (quantified in (F)).

**Figure S6:**
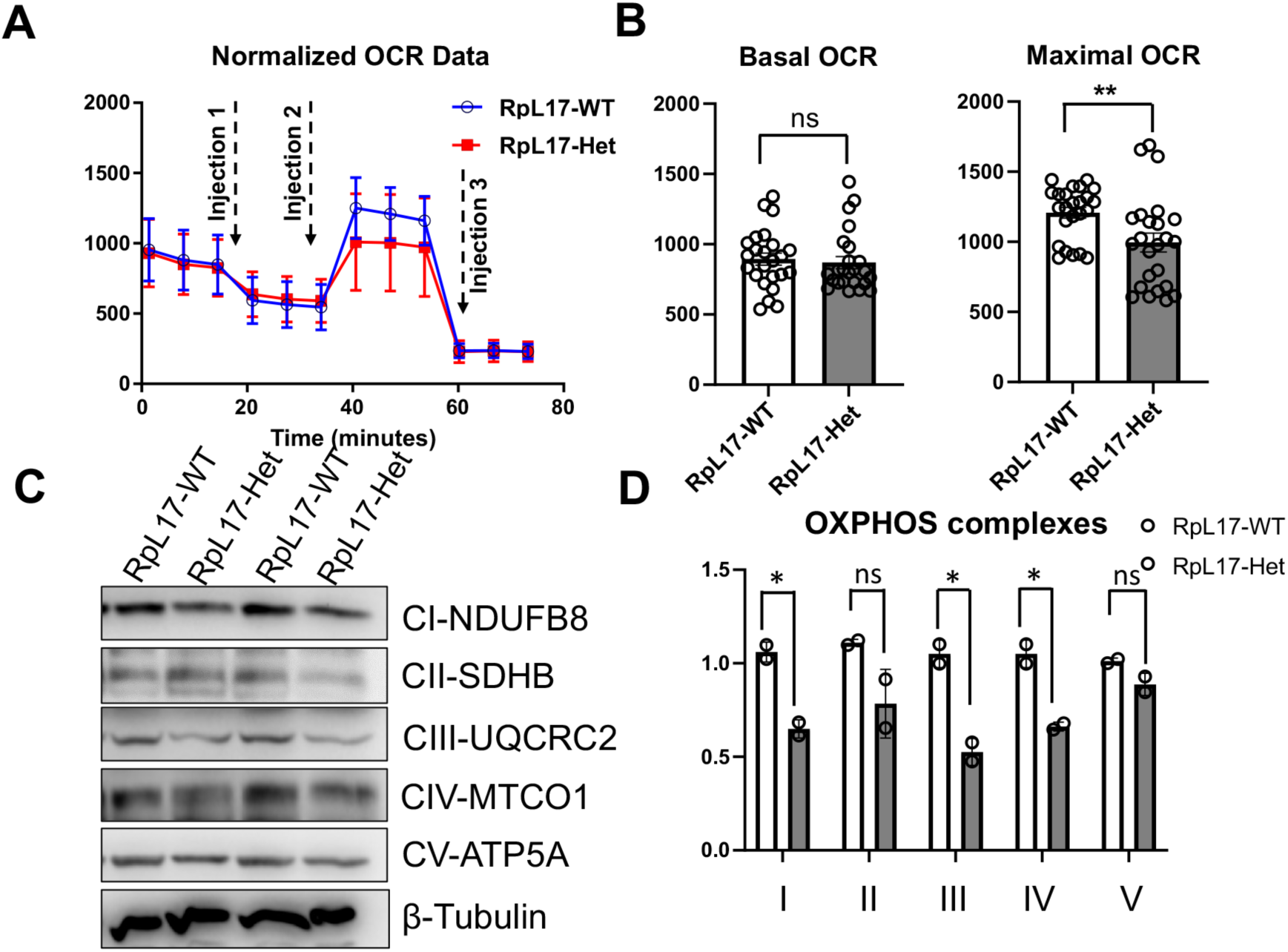
Decreased RpL17 in EC leads to mitochondrial dysfunction. (A) Seahorse extracellular flux analysis of mitochondrial (mt) function in RpL17-WT vs. RpL17-Het MLMEC identified decreased mt respiration (≈ 16%) in RpL17-Het compared to RpL17-WT. Injections 1: 1µM oligomycin; 2: 2µM FCCP; and 3: 1µM (rotenone + antimycin A). (B) Quantification of the oxygen consumption rate (OCR) in (A) revealed no change in basal OCR (B), but a significant decrease in maximal (B) OCR in RpL17-Het compared to RpL17-WT (Average of n=8). (C) Decreased expression of OXPHOS complex proteins (CI-NDUFB8**, CII-SDHB**, CIII-UQCRC2**, CIV-MTCO1, and CV-ATP5A) in RpL17-Het compared to RpL17-WT; quantified in (D)(n=2; p< 0.05).

